# Sequence-based Drug-Target Binding Site Pre-training Enables Cryptic Pocket Detection and Improves Binding Affinity and Kinetics Prediction

**DOI:** 10.1101/2025.01.14.633076

**Authors:** Shuo Zhang, Li Xie, Daniel Tiourine, Lei Xie

## Abstract

Predicting protein-ligand binding characteristics, such as affinity and kinetics, is critical for accelerating drug discovery. However, many existing computational methods face key limitations, including insufficient integration of comprehensive databases, inadequate representation of protein structural dynamics, and incomplete modeling of microscale protein-ligand interactions. To address these challenges, we introduce ProMoNet, a sequence-based pre-training and fine-tuning framework to enhance the prediction of protein-ligand binding characteristics. ProMoNet leverages protein and molecular foundation models to expand data coverage and enhance diversity. It also introduces a pre-training strategy based on protein-ligand binding site prediction, which bridges protein- and ligand-level representations to support downstream prediction tasks involving protein-ligand complexes. Our pre-training module effectively models microscale protein-ligand interactions and captures the dynamic nature of proteins, including binding site crypticity, without relying on 3-dimensional structural inputs. Notably, this module surpasses or matches state-of-the-art structure-based methods in identifying exposed and cryptic binding sites while maintaining high efficiency. Our fine-tuning module then efficiently transfers the pre-trained knowledge to downstream tasks such as binding affinity and binding kinetics prediction, achieving superior performance. The combination of ProMoNet’s strong performance and demonstrated efficiency across multiple tasks highlights its potential for broad applications in drug discovery.

**Scientific Contribution:** We propose ProMoNet, a sequence-based pre-training and fine-tuning framework for protein-ligand binding characteristic prediction, where protein-ligand binding site prediction is introduced as a pre-training strategy to bridge independent protein- and ligand-level representations for downstream complex-level tasks. We design two dedicated modules, including a pre-training module that models microscale protein-ligand interactions and captures protein dynamics, as well as a fine-tuning module that efficiently integrates the pre-trained representations for downstream tasks. Even compared to structure-based methods, ProMoNet matches state-of-the-art performance in exposed and cryptic binding site identification and delivers superior results in binding affinity and kinetics prediction, making it a promising tool for drug discovery.

## 1 Introduction

Characterizing the binding process between proteins and small-molecule ligands is a critical step in drug discovery. These binding characteristics encompass structural properties like binding sites and poses [1, 2], thermodynamic properties including equilibrium constants *K*_i_, *K*_d_ [3], and kinetic properties such as rate constants *k*_on_, *k*_off_ [4]. Together, these properties provide essential insights into protein-ligand interactions, facilitating more effective compound screening [5]. To accelerate the prediction of protein-ligand binding characteristics and reduce associated costs, various computational approaches have been developed, demonstrating promising performance [6].

Despite significant progress in predicting protein-ligand binding characteristics, existing methods still face several critical limitations. First, *the datasets used for training models are often limited and biased*. When experimental data for a specific binding characteristic is scarce, the performance of the trained models tends to suffer [7]. Additionally, while binding characteristics are closely interrelated [5], existing models often focus exclusively on the dataset for the characteristic being predicted and fail to leverage relevant information from datasets corresponding to other characteristics. This neglect of cross-characteristic information further limits their ability to generalize and accurately predict binding characteristics. Second, *the dynamic nature of protein structures is often overlooked or inefficiently captured*. Many structure-based methods rely solely on rigid 3-dimensional (3D) protein structures, failing to account for the dynamic interactions during binding [8–10]. Although some approaches address dynamics, they often rely on resource-intensive molecular dynamics (MD) simulations [11–15] or time-consuming diffusion-based generative modeling [16, 17], making them impractical for high-throughput compound screening. Third, *many models rely solely on protein- and molecule-level representations, neglecting their microscale interactions* [18–20]. These interactions are essential for capturing the fundamental mechanisms of binding and are critical for accurately predicting characteristics of protein-ligand binding characteristics [21–23].

To address the aforementioned limitations, we present **ProMoNet**, a sequence-based pre-training and fine-tuning framework designed to enhance protein-ligand binding characteristic prediction. ProMoNet bridges the representations from independent protein and molecular foundation models, leveraging recent advancements in foundation models [24–26] and their pre-training and fine-tuning strategies [27, 28]. ProMoNet consists of two key modules. The **pre-training module**, ProMoSite, learns microscale protein-ligand interactions and captures dynamic structural information, including binding site crypticity by training on binding site annotations in protein-ligand complexes. The **fine-tuning module**, ProMoBind, integrates ProMoSite’s pre-trained capabilities to be fine-tuned for predicting protein-ligand binding characteristics, including binding affinity and kinetics. Although previous studies have explored leveraging protein–ligand interaction information to improve binding affinity prediction [29–31], our work introduces binding site prediction as a pre-training strategy to bridge independent protein and molecular foundation models, enabling a unified framework for diverse binding characteristic prediction tasks.

To comprehensively evaluate ProMoNet, we assess its two modules, ProMoSite and ProMoBind, on diverse protein–ligand binding prediction tasks. ProMoSite is tested on binding site prediction during pre-training, including both *exposed* sites on the protein surface and *cryptic* sites that are not readily observable in *apo* protein structures, but appear through ligand-induced conformational changes [14, 32]. Using only sequence data, it matches or surpasses state-of-the-art methods that depend on 3D structures and achieves the best results on cryptic site prediction, highlighting its ability to capture protein dynamics from sequence alone. ProMoBind is evaluated on binding affinity prediction and binding kinetics prediction. ProMoBind uses only sequence data and leverages the pre-trained ProMoSite, achieving high efficiency and outperforming state-of-the-art baselines across both tasks. These results demonstrate ProMoNet’s potential as a robust and efficient sequence-based framework for high-throughput compound screening and broader applications in drug discovery.

## 2 Results and discussion

### 2.1 Overview of ProMoNet

The overall architecture of ProMoNet, including its two key modules, ProMoSite and ProMoBind, is illustrated in Figure 1. ProMoSite (Fig. 1a) serves as the pre-training module that connects independent protein and molecular foundation models via pre-training on binding site databases [33, 34]. It predicts whether an amino acid residue physically interacts with a ligand (i.e., binding site residue) and is evaluated for both exposed and cryptic binding site identification. ProMoBind (Fig. 1b), the fine-tuning module, builds on the pre-trained ProMoSite to be efficiently fine-tuned on downstream protein-ligand binding characteristics prediction tasks, such as binding affinity and kinetics prediction. The pre-training and fine-tuning workflow of ProMoNet is illustrated in Fig. 1c, and the experimental setup for evaluating ProMoNet’s modules is shown in Fig. 1d. The details of ProMoNet are described in the following paragraphs.

**Fig. 1.**
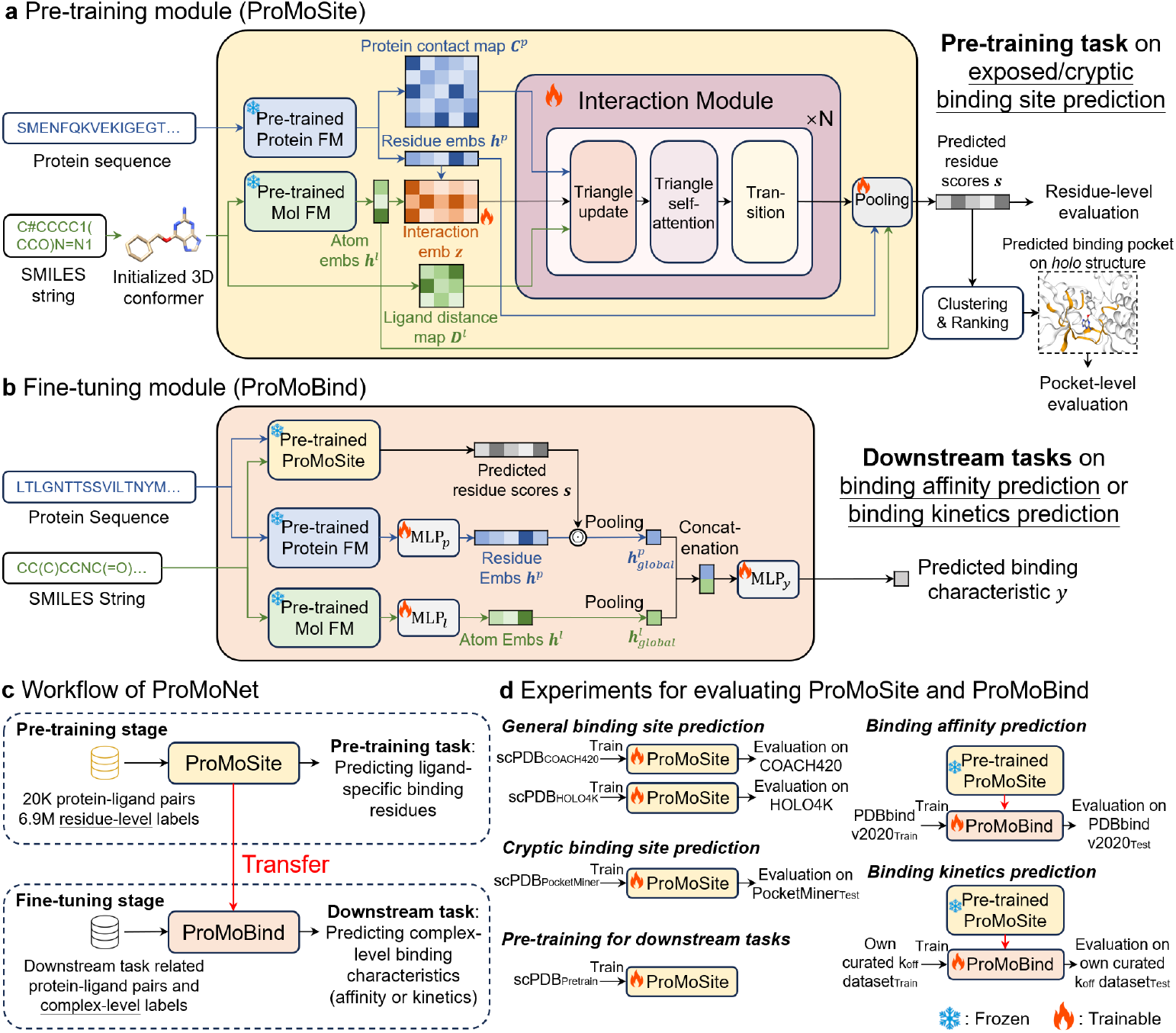
Overview of ProMoNet. ProMoNet consists of two key modules: ProMoSite, the pretraining module, and ProMoBind, the fine-tuning module. (**a**) ProMoSite employs foundation models (FMs) to extract initial embeddings from protein sequences and ligand SMILES strings. We then define protein-ligand microscale interaction embeddings and iteratively update them through our interaction module, constrained by protein contact maps and ligand distance maps. The updated embeddings are used to compute residue-level scores for binding site prediction. (**b**) ProMoBind incorporates the pre-trained ProMoSite to predict complex-level protein-ligand binding characteristics with multiple trainable multi-layer perceptions (MLPs). This efficient fine-tuning and inference design makes ProMoBind scalable and suitable for high-throughput compound screening. (**c**) Pretraining and fine-tuning workflow of ProMoNet. (**d**) Experimental setup for evaluating ProMoSite and ProMoBind. The snowflake icon (“Frozen”) indicates components whose parameters are kept fixed during training, while the flame icon (“Trainable”) indicates components with trainable parameters.

ProMoSite (Fig. 1a) takes advantage of the transferability of foundation models, which are pre-trained on extensive databases to increase data size and diversity [24]. In detail, we adopt ESM-2 [35] as the protein foundation model and use Uni-Mol [36] as the molecular foundation model. Both ESM-2 and Uni-Mol are pre-trained and the parameters are kept frozen during the pre-training of ProMoSite. To represent protein-ligand pairs, ProMoSite takes the protein sequences and the ligand SMILES strings [37] as input. The protein sequences are fed into ESM-2 to get residue embeddings and predict protein contact maps, while the SMILES strings are used to initialize 3D conformers via RDKit [38] to compute distance maps and as the input of Uni-Mol to get atom embeddings. Based on each pair of residue embedding and atom embedding, ProMoSite forms a protein-ligand interaction embedding to model the microscale interactions between protein residues and ligand atoms. Furthermore, the interaction embedding is updated via ProMoSite’s interaction module which utilizes protein contact maps and ligand distance maps as triangle inequality constraints. Unlike previous works [9, 39] that aim to satisfy the constraints solely on distances, ProMoSite avoids using rigid 3D structure-based distance maps for proteins while using contact maps instead. As shown in prior studies [40, 41], protein contact maps can capture information related to multistate protein conformations. By using contact maps, our model can implicitly leverage dynamic structural information associated with multistate protein conformations when modeling microscale protein-ligand interactions. Besides, ProMoSite’s interaction module uses a self-attention mechanism [39, 42] to model dependencies among residue–atom interactions, allowing each interaction to be updated by attending to other residue–atom interaction pairs. It accounts for physical and chemical effects such as steric constraints (excluded-volume, van der Waals interactions) and saturation effects [9] for more effective modeling. The updated interaction embedding from the interaction module as well as the initial residue and atom embeddings are fed into the pooling module to quantify a score between each residue and the entire ligand, where a higher score indicates a greater likelihood that the residue belongs to a binding site of the given ligand. Such scores can be directly used for residue-level metrics (e.g. ROC-AUC, PR-AUC) when evaluating ProMoSite for binding site identification. To additionally evaluate the identified binding site in 3D space, we post-process the scored residues with clustering and ranking. The filtered residues are mapped onto known *holo* structures to compute pocket-level metrics like DCA (distance from the center of the predicted binding pocket to the closest ligand heavy atom). To pre-train ProMoSite, we leverage protein-ligand binding site annotations from large-scale, open-source databases [33, 34] as signals of microscale interactions, enabling ProMoSite to effectively learn and predict them.

ProMoBind (Fig. 1b) adopts the same input formats as ProMoSite to predict the binding characteristic of a given protein–ligand pair, while still utilizing ESM-2 and Uni-Mol to extract protein residue embeddings and ligand atom embeddings. To achieve efficiency in fine-tuning, we freeze all parameters in these pre-trained foundation models and adapt trainable multi-layer perceptions (MLP) after the extracted embeddings to improve generalization and transferring ability on down-stream tasks [28, 43]. Then ProMoBind incorporates the predicted residue scores from pre-trained ProMoSite by taking an element-wise product with the residue embeddings, which can apply weighting to these embeddings according to the protein-ligand interaction likelihoods that exist in predicted scores. Moreover, the resulting residue embeddings and atom embeddings are pooled to get corresponding global embeddings of protein and ligand. Finally, ProMoBind concatenates the pair of global embeddings and uses a trainable MLP-based prediction head to predict the desired protein-ligand binding characteristic. In ProMoBind, only the parameters of three MLP modules are fine-tuned, and the foundation models only require a single forward pass during inference. Such efficiency of ProMoBind in training and inference is invaluable for resource-intensive tasks like high-throughput compound screening.

### 2.2 ProMoSite excels in predicting general binding sites and residue-level non-covalent interaction sites

To evaluate ProMoSite’s performance for identifying *general binding sites*, we used the scPDB v2017 database [33] for training and cross-validation, and benchmark datasets including COACH420 and HOLO4K [8] for testing. To avoid data leakage, we adopted the same strategy and data split as in [44], ensuring that the proteins in the training and test sets had neither similar global structures nor similar binding sites. For detailed dataset statistics and processing steps, please refer to the Dataset subsection in the Methods section.

For a comprehensive evaluation, we compared ProMoSite against two types of state-of-the-art methods, including 3D structure-based (Fpocket [45], Deepsite [46], P2Rank [8], Kalasanty [47], PUResNet [48], DeepPocket [44]) and 1D sequence-based methods (HoTS [30], DeepProSite [49], Pseq2Sites [50]). The 3D structure-based methods require 3D protein coordinate information as input. Their predicted binding sites are evaluated in 3D space using metrics such as DCA, where predictions with DCAs *<* 4Å are considered successful [8, 44]. Based on the DCA criterion, we evaluated the models’ ranking capability by measuring the success rates for the top n-ranked predictions, following [8, 44], where n is the number of annotated binding sites for a given protein. For 1D sequence-based methods, only protein sequences are required as input. Since these methods directly predict the probabilities of protein residues belonging to binding sites, binary classification metrics are typically used [49]. Specifically, we used the area under the receiver operating characteristic curve (ROC-AUC) and the area under the precision-recall curve (PR-AUC) for comparison.

As shown in Fig. 2a, ProMoSite outperforms all 3D structure-based baselines, achieving an averaged Top-n success rate of 68.30% ± 0.99 on COACH420 and 73.46% ± 0.65 on HOLO4K, respectively. These results demonstrate that ProMoSite, which requires only protein sequences as input, effectively incorporates 3D structural information that emerged from the pre-trained ESM-2 model to identify ligand binding sites in 3D space, including on both single-chain and multi-chain proteins. The ablation study results shown in Supplementary Table 1 further validate the effectiveness of ProMoSite’s interaction module and pooling module, which play critical roles in modeling protein-ligand interactions.

**Fig. 2.**
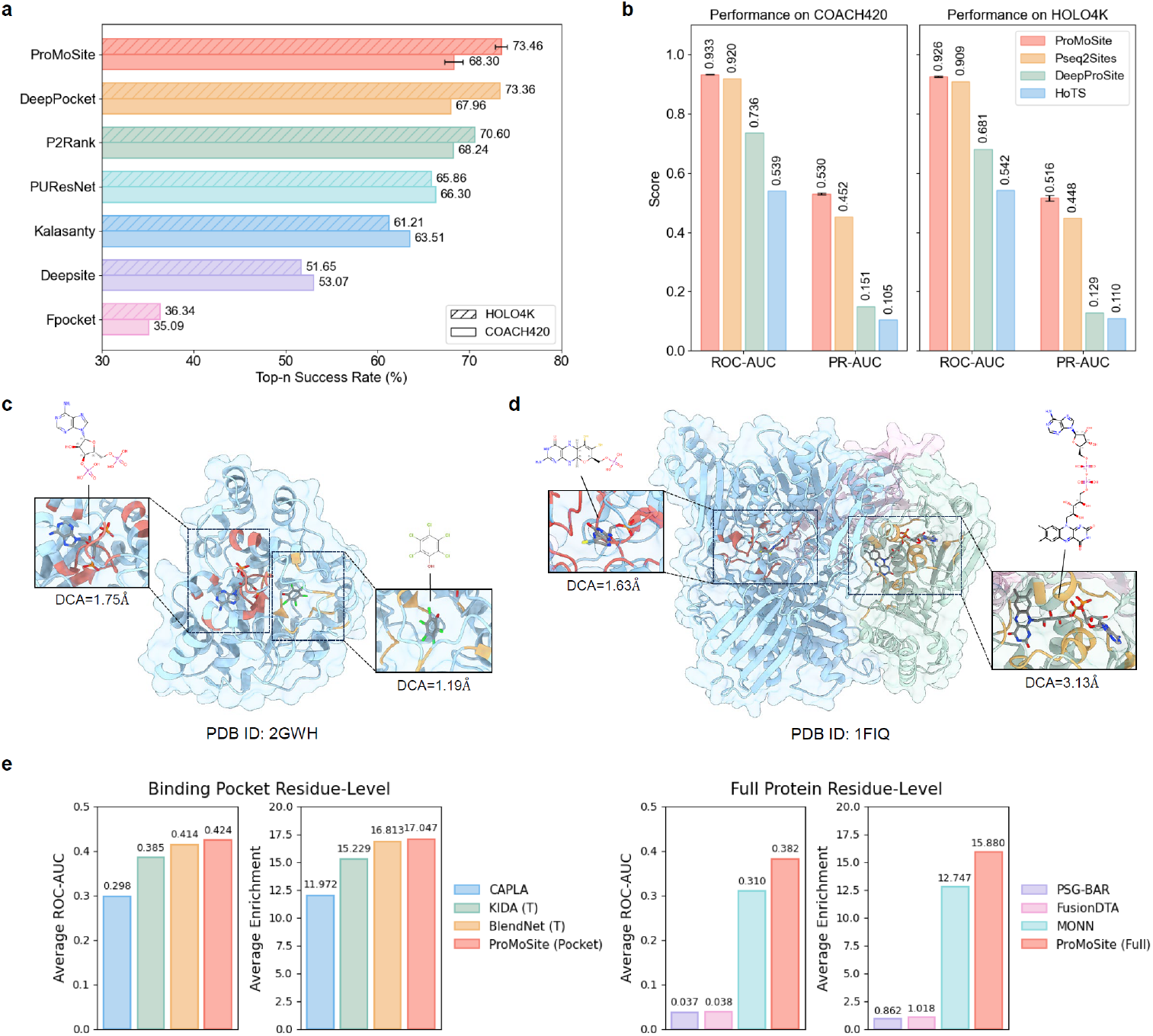
Comparison of model performance in predicting general binding sites. (**a**) ProMoSite outperforms all 3D-based baselines on COACH420 and HOLO4K, achieving high Top-n success rates for identifying ligand binding sites in 3D space using only sequence data as input. (**b**) ProMoSite outperforms sequence-based baselines on COACH420 and HOLO4K by leveraging pretrained protein and molecular foundation models and incorporating geometric constraints via contact maps and distance maps. (**c, d**) Representative examples of ligand-specific binding site predictions by ProMoSite on proteins with multiple ligands, demonstrating its capability to accurately predict specific binding sites for each ligand. (**e**) ProMoSite outperforms baselines on MISBD for residue-level non-covalent interaction sites detection.

As shown in Fig. 2b and Supplementary Fig. 1, when compared to 1D sequence-based methods, ProMoSite consistently outperforms them on COACH420 (ROC-AUC = 0.933 ± 0.001; PR-AUC = 0.530 ± 0.003) and HOLO4K (ROC-AUC = 0.926 ± 0.001; PR-AUC = 0.516 ± 0.009). The superior performance of ProMoSite demonstrates the effectiveness of our approach. Compared with HoTS, which uses 1D-CNN [51] and Morgan fingerprints [52] to extract protein and molecular representations, respectively, ProMoSite leverages pre-trained protein and molecular foundation models to improve model generalization. Although HoTS explicitly models ligand-protein interactions using transformer blocks [42], it lacks geometric constraints in protein-ligand binding. In contrast, ProMoSite incorporates protein contact maps and ligand distance maps as triangle inequality constraints to model microscale ligand-protein interactions, achieving consistently better performance. Compared with DeepProSite, which uses ESMFold-predicted 3D protein structures and incorporates a pre-trained protein language model for sequence embeddings, ProMoSite eliminates the need for 3D protein structures. Instead, it uses predicted protein contact maps to capture the dynamic structural information of multistate protein conformations. Compared with Pseq2Sites, which does not consider ligand information in binding site detection, ProMoSite explicitly models microscale ligand-protein interactions and can produce *ligand-specific* binding site predictions when a given protein binds to multiple different ligands at different sites.

While most methods in our comparison only identify unspecific binding sites, a reliable prediction method should be able to identify the specific binding site for a query ligand [53]. Among the baselines in our experiments, only HoTS and DeepProSite can predict ligand-specific binding sites. However, DeepProSite requires retraining for each ligand type, limiting its practical usability. In contrast, both HoTS and ProMoSite directly take ligand information as input, enabling the same model to generalize across any ligand structure. To demonstrate ProMoSite’s capability of identifying ligand-specific binding sites, we provide representative visualizations (Fig. 2c, d) of its predictions in scenarios where a protein binds to multiple ligands at different sites. Fig. 2c shows a prediction example from COACH420 on human sulfotransferase SULT1C2 in complex with PAP and pentachlorophenol (PDB ID: 2GWH). ProMoSite accurately identifies the two specific binding sites corresponding to their ligands. Similarly, Fig. 2d presents a prediction example from HOLO4K on xanthine oxidase in complex with FAD and molybdopterin cofactor (PDB ID: 1FIQ). In this multi-chain protein example, ProMoSite accurately identifies ligand-specific binding sites across different chains, further demonstrating its effectiveness.

Although ProMoSite is trained using residue-level binding site annotations and directly predicts residue-level scores between 0 and 1 associated with binding sites, we further investigate whether these scores can capture non-covalent interactions at residue-level and provide interpretability as pursued in previous works [29, 31]. We evaluated ProMoSite on the Merged Interaction Sites Benchmark Dataset (MISBD) [31] and compared it with the baselines in [31], including PSG-BAR [54], FusionDTA [55], CAPLA [56], MONN [29], KIDA [57], and BlendNet [31]. Since ProMoSite predicts residue-level scores, we use these scores together with the residue-level non-covalent interaction labels in MISBD to compute the area under the receiver operating characteristic curve (ROC-AUC) and enrichment scores following [31]. Moreover, we found that CAPLA, KIDA, and BlendNet only produce residue scores in protein pocket regions. Therefore, for the evaluation in [31], the non-pocket residues are padded with 0 to calculate the metrics based on the full protein sequence. For fair comparison, we use the same pocket annotations provided by [31] to mask the non-pocket residues with 0 for our predicted residue scores when computing the metrics. We refer to this prediction as ProMoSite (Pocket). In contrast, PSG-BAR, FusionDTA, and MONN can produce residue scores for the full protein sequence without any padding. Therefore, for fair comparison with these models, we directly use our raw predicted residue scores when computing the metrics. We refer to this prediction as ProMoSite (Full). As shown in Fig. 2e, ProMoSite variants consistently outperform the corresponding baselines on all metrics when detecting residue-level non-covalent interaction sites in binding pockets or across full protein sequences, demonstrating its strong capacity to capture residue-level protein–ligand non-covalent interaction patterns from binding residue annotations. These results further suggest that the residue-level scores produced by ProMoSite reflect biologically relevant interaction patterns, as residues assigned higher scores are more frequently associated with ligand-contacting regions and non-covalent interaction sites. Together, these findings support the residue-level interpretability of ProMoSite.

### 2.3 ProMoSite matches state-of-the-art performance in cryptic binding site detection with high efficiency

Following our analysis of general binding site prediction, we also evaluate ProMoSite’s performance in identifying *cryptic binding sites*, which are ligand-binding pockets that are not apparent in *apo* protein structures but emerge upon ligand binding or protein conformational changes [14, 32, 58]. Accurately detecting these sites is crucial for understanding allosteric regulation and screening potential drugs for proteins currently considered undruggable [32]. For this task, we trained ProMoSite using the scPDB v2017 database [33] and evaluated it on the PocketMiner validation and test sets [14], which are specifically designed for cryptic binding site prediction. Details of these datasets are provided in the Methods section.

We compared ProMoSite with several state-of-the-art models. CryptoSite [59] and PocketMiner [14] require 3D protein structures obtained from MD simulations and directly output residue-level probabilities for cryptic binding sites. NeuralPLexer [16] and DynamicBind [17] are diffusion-based generative models that can reveal cryptic binding sites by generating 3D ligand-protein binding complexes. We evaluated two variants of NeuralPLexer based on the type of protein-related input: NeuralPLexer-fasta, which uses protein sequences, and NeuralPLexer-AF2, which utilizes AlphaFold2 [39]-predicted 3D protein structures. Additionally, we included P2Rank, which requires 3D protein structures as input but focuses on general binding site prediction. To use P2Rank for cryptic binding site prediction, we provide 3D *apo* protein structures as input, which haven’t formed the potential cryptic pockets already. While most of the baselines rely on 3D protein structures, ProMoSite is sequence-based and trained on general binding site databases without using MD simulation data or protein conformations related to cryptic binding sites.

For evaluation, we adopted different metrics depending on the model outputs. For CryptoSite and PocketMiner, we used ROC-AUC and PR-AUC following [14], as they output residue-level probabilities. For NeuralPLexer, DynamicBind, and P2Rank, whose predictions are either generated 3D complexes or 3D binding sites, we used the DCA metric. Additionally, for NeuralPLexer and DynamicBind, residues within 7Å of any ligand heavy atom in the generated binding complex were considered as cryptic binding site residues. To ensure consistency, all predicted residues were mapped onto experimental *holo* protein structures when calculating DCA. More details of the experimental settings are provided in the Methods section.

Fig. 3a presents a comparison of the ROC and PR curves for ProMoSite, CryptoSite, and PocketMiner on the test set. The ROC curves demonstrate that ProMoSite outperforms CryptoSite and PocketMiner, achieving a higher ROC-AUC of 0.91 ± 0.01. Similarly, the PR curves show that ProMoSite maintains superior performance across a broader range of recall levels, achieving a higher PR-AUC of 0.84 ± 0.01. These results highlight ProMoSite’s effectiveness. Even when trained solely on a general binding site dataset and without requiring MD-derived data or 3D protein structures, it surpasses state-of-the-art models that depend on 3D structures obtained from MD simulations.

**Fig. 3.**
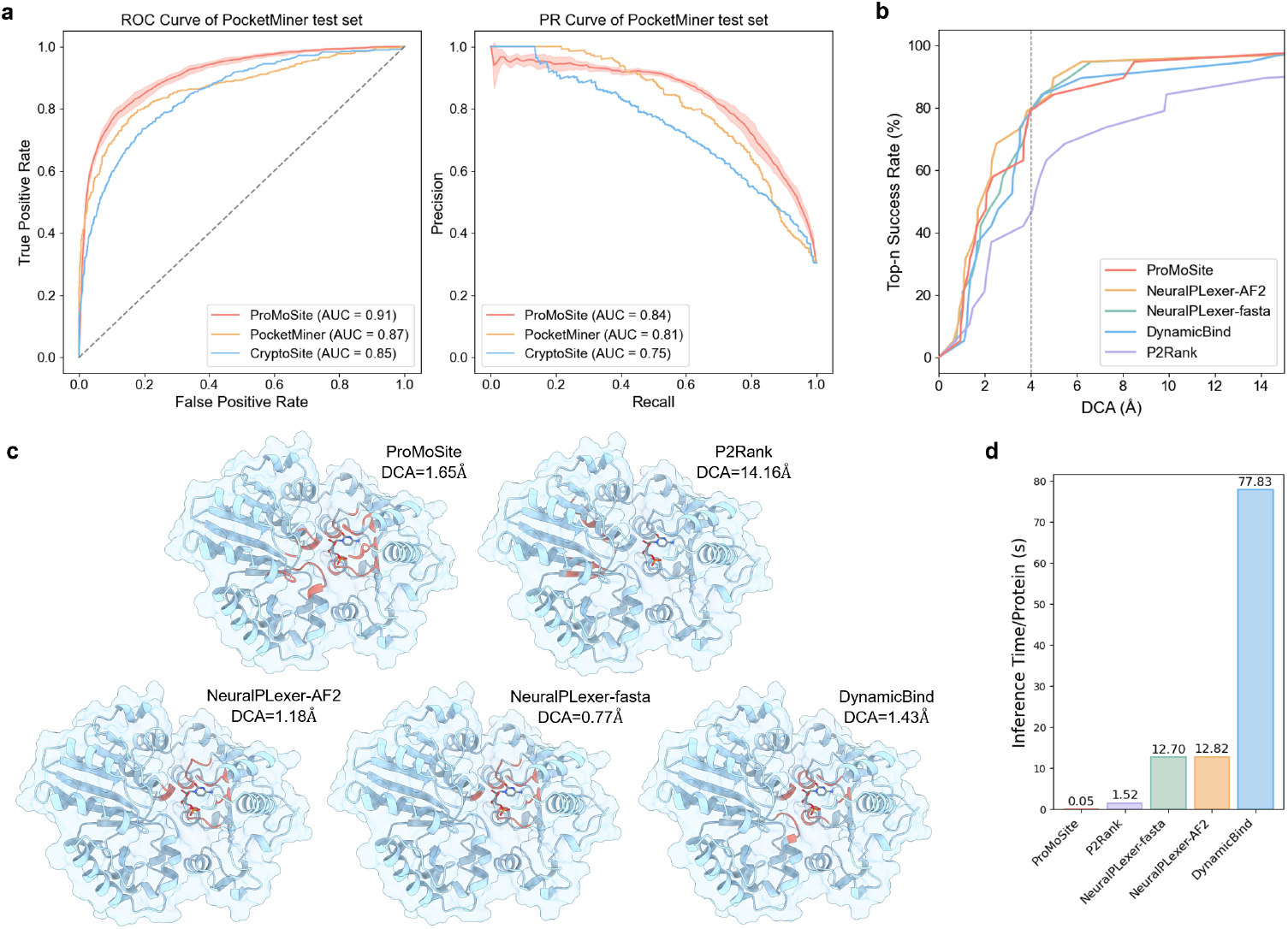
Comparison of model performance in predicting cryptic binding sites. (**a**) ROC and PR curves comparing ProMoSite with CryptoSite and PocketMiner on the test set. ProMoSite achieves the highest ROC-AUC and PR-AUC, demonstrating superior performance even without requiring MD-derived data or 3D protein structures. (**b**) Top-n success rates as a function of DCA values for ProMoSite and baselines. At a 4.0Å threshold, ProMoSite matches NeuralPLexer-AF2 and NeuralPLexer-fasta while outperforming DynamicBind and P2Rank. (**c**) Visualization of cryptic binding site predictions for *α*-2,3-sialyltransferase using the *apo* structure (PDB ID: 4V38), mapped onto the experimental *holo* structure (PDB ID: 4V3B). ProMoSite, NeuralPLexer-AF2, NeuralPLexer-fasta, and DynamicBind accurately identify the cryptic binding site, whereas P2Rank fails due to its reliance on rigid protein structures. (**d**) Inference speed comparison among models. ProMoSite achieves the fastest inference time (0.05s/protein) while maintaining cryptic binding site identification performance comparable to generative models.

In Fig. 3b, we compare the Top-n success rate as a function of DCA values for ProMoSite and relevant baselines. The detailed DCA values are listed in Supplementary Table 3. ProMoSite achieves performance comparable to the current state-of-the-art generative models NeuralPLexer-AF2, NeuralPLexer-fasta, and DynamicBind, while outperforming P2Rank. Specifically, at the commonly used threshold of 4.0Å, ProMoSite, NeuralPLexer-AF2, and NeuralPLexer-fasta all achieve a success rate of 78.9%, while DynamicBind achieves 73.7%. However, P2Rank, designed for general binding site prediction, only achieves a success rate of 42.1%.

As a representative example, Fig. 3c visualizes the cryptic binding site predictions of these models when provided with the *apo* structure or sequence of *α*-2,3-sialyltransferase (PDB ID: 4V38), mapped onto the corresponding experimental *holo* structure (PDB ID: 4V3B). The visualizations show that all models except P2Rank accurately identify the cryptic binding site, which emerges through secondary structure motion. P2Rank fails to detect this site that is absent in the *apo* structure, as it treats protein structures as rigid and cannot capture protein dynamics. In contrast, when modeling microscale protein-ligand interactions, ProMoSite uses predicted protein contact map as geometric constraints, which can capture the dynamic structural information of multistate protein conformations [40, 41]. From a biological perspective, the ability to recover such cryptic pockets is important because these sites are often associated with conformationally regulated or allosteric binding events that are difficult to identify from rigid apo structures alone. This result further suggests that ProMoSite captures sequence-derived signals related to conformational adaptability that are relevant to cryptic pocket formation.

Further analysis in Fig. 3d compares the inference speed of these models. While the generative models NeuralPLexer-AF2, NeuralPLexer-fasta, and DynamicBind exhibit strong cryptic binding site identification performance, their inference times per protein are considerably slower due to their reliance on diffusion models [60] when generating 3D structures (NeuralPLexer-AF2: 12.82s, NeuralPLexer-fasta: 12.70s, DynamicBind: 77.83s). P2Rank, with an inference time of 1.52s per protein, is faster but delivers much worse performance in cryptic binding site identification. In contrast, ProMoSite achieves cryptic binding site identification performance on par with these structure-based state-of-the-art models (Fig. 3b) while offering the fastest inference speed at just 0.05s per protein. This combination of speed and accuracy demonstrates ProMoSite’s potential for large-scale virtual screening and drug discovery applications.

### 2.4 ProMoBind outperforms baselines in binding affinity prediction

Following the evaluation of ProMoSite’s superior performance in predicting general and cryptic binding sites, we now assess its downstream application in predicting binding characteristics using ProMoBind, which leverages the outputs from pre-trained ProMoSite. Specifically, we focus on binding affinity, one of the most critical and representative binding characteristics [6]. To evaluate ProMoBind’s performance, we conducted experiments on the commonly used PDBbind v2020 [75] benchmark. We used four metrics: Root Mean Square Error (RMSE), Mean Absolute Error (MAE),

Pearson correlation coefficient, and Spearman correlation coefficient. We also evaluated our model on eight test datasets from multiple sources (PDBbind [75], CASF [76], CSAR [77], and Astex [78]) to enable a comprehensive comparison. ProMoSite was pre-trained on the scPDB v2017 database [33] to predict residue scores, which were then integrated into ProMoBind. To assess the impact of these scores, we developed a variant of ProMoBind for comparison, named ProMoBind-Base, which excludes ProMoSite predicted scores for comparison.

For the experiments on PDBbind v2020, we first applied a time-based split following [9, 23], where data points published before 2019 were used for training and validation, and those published during or after 2019 formed the test set. Each sample in PDBbind v2020 includes an experimentally determined 3D protein-ligand complex and its binding affinity, expressed in negative log-transformed dissociation (p*K*_d_) or inhibition (p*K*_i_) constant values. Accordingly, we compared ProMoBind against state-of-the-art baselines that utilize different types of protein-related inputs, including 3D protein-ligand complex structures (Pafnucy [61], OnionNet [62], IGN [63], and SIGN [21]), 3D *holo* protein structures (SMINA [64], GNINA [65], dMaSIF [66], TankBind [9], DynamicBind [17], and Interformer [67]), protein sequences (TransformerCPI [68], MolTrans [69], GraphDTA [19], and DrugBAN [70]), protein sequences with predicted protein contact maps (DGraphDTA [71], WGNN-DTA [72], STAMP-DPI [73], and PSICHIC [23]), and protein sequences with predicted 3D complex structures (Boltz-2 [74]). The results of all models are summarized in Table 1, where ProMoBind achieves the highest Pearson and Spearman correlation coefficients and the second-lowest RMSE. Overall, it outperforms all baselines that use various types of protein-related inputs. Even ProMoBind-Base, which excludes the ProMoSite predicted scores, achieves the second-best scores on Pearson and Spearman correlation coefficients. This highlights the superiority and robustness of the protein and molecular foundation models pre-trained on diverse datasets, as well as the effectiveness of our approach in leveraging these models. Among the baselines, although DynamicBind achieves the lowest RMSE and MAE, its correlation metrics are substantially lower than those of our models. Moreover, as illustrated in Fig. 3d, DynamicBind is computationally inefficient, requiring 77.83 seconds/protein–ligand pair during inference because it must first generate 3D binding complex structures from separate protein and ligand structures before predicting affinity. In contrast, ProMoBind is the fastest model among all baselines, requiring only 0.0012 seconds/protein–ligand pair. Compared to ProMoBind-Base, ProMoBind achieves consistent performance improvements (Table 1 and Supplementary Table 4), indicating that the incorporated ProMoSite predicted scores effectively capture microscale protein-ligand interaction information in our pre-training, thereby enhancing the model’s ability to predict binding affinity more accurately. These gains further suggest that residue-level binding-site priors provide biologically relevant cues for complex-level prediction by highlighting protein regions that are more likely to mediate ligand recognition and affinity.

**Table 1.** Comparison of model performance in predicting binding affinity on PDBbind v2020 using time-based split. The protein-related input type for each model is indicated. ‘1D Seq → 2D Map’ denotes models that use both 1D protein sequences and their predicted 2D contact maps as input. ‘1D Seq → 3D Complex’ denotes the model requires 1D protein sequences to generate 3D complex for prediction. The best results are highlighted in bold, and the second-best results are underlined.

| Methods | Protein-related Input | RMSE ↓ | MAE ↓ | Pearson ↑ | Spearman ↑ |
| --- | --- | --- | --- | --- | --- |
| Pafnucy [61] | 3D Complex | 1.435 | 1.144 | 0.635 | 0.587 |
| OnionNet [62] | 3D Complex | 1.403 | 1.103 | 0.648 | 0.602 |
| IGN [63] | 3D Complex | 1.404 | 1.116 | 0.662 | 0.638 |
| SIGN [21] | 3D Complex | 1.373 | 1.086 | 0.685 | 0.656 |
| SMINA [64] | 3D Struct | 1.466 | 1.161 | 0.665 | 0.663 |
| GNINA [65] | 3D Struct | 1.740 | 1.413 | 0.495 | 0.494 |
| dMaSIF [66] | 3D Struct | 1.450 | 1.136 | 0.629 | 0.588 |
| TankBind [9] | 3D Struct | 1.345 | 1.060 | 0.718 | 0.689 |
| DynamicBind [17] | 3D Struct | **1.253** | **1.013** | 0.689 | 0.656 |
| Interformer [67] | 3D Struct | - | - | 0.659 | - |
| TransformerCPI [68] | 1D Seq | 1.493 | 1.201 | 0.604 | 0.551 |
| MolTrans [69] | 1D Seq | 1.599 | 1.271 | 0.539 | 0.474 |
| GraphDTA [19] | 1D Seq | 1.564 | 1.223 | 0.612 | 0.570 |
| DrugBAN [70] | 1D Seq | 1.480 | 1.159 | 0.657 | 0.612 |
| DGraphDTA [71] | 1D Seq → 2D Map | 1.493 | 1.201 | 0.604 | 0.551 |
| WGNN-DTA [72] | 1D Seq → 2D Map | 1.501 | 1.196 | 0.605 | 0.562 |
| STAMP-DPI [73] | 1D Seq → 2D Map | 1.503 | 1.176 | 0.653 | 0.601 |
| PSICHIC [23] | 1D Seq → 2D Map | 1.314 | <u>1.015</u> | 0.710 | 0.686 |
| Boltz-2 [74] | 1D Seq → 3D Complex | - | - | 0.609 | 0.600 |
| **ProMoBind-Base** | 1D Seq | 1.290 | 1.041 | <u>0.725</u> | <u>0.694</u> |
| **ProMoBind** | 1D Seq → 2D Map | <u>1.268</u> | 1.020 | **0.738** | **0.707** |

To evaluate the generalization power of ProMoBind, we further applied a protein sequence identity-based data split on PDBbind v2020 by removing proteins from our time-based training and validation sets that have more than 80% sequence identity with any protein in the test set. The overall best-performing baseline, PSICHIC, along with our ProMoBind and ProMoBind-Base, were included for comparison. As shown in Supplementary Tables 5 and 6, ProMoBind consistently achieves the best performance across all four evaluation metrics, demonstrating its strong generalization ability. Furthermore, ProMoBind shows consistent improvements over ProMoBind-Base, highlighting the effectiveness of incorporating the pre-trained ProMoSite.

Besides evaluating on PDBbind v2020, we conducted the same binding affinity prediction experiment as in [31] to compare with more baselines and evaluate on more diverse datasets. We retrained ProMoBind using the same training set and data split as in [31], and evaluated the model on eight test sets, including one internal test set and seven external datasets from PDBbind [75], CASF [76], CSAR [77], and Astex [78]. We report the detailed Pearson correlation coefficient (PCC) of each model on the eight test sets in Supplementary Fig. 2a, and report the averaged ranking of each model based on PCC across the eight test sets in Supplementary Fig. 2b. The PCC ranking score results show that ProMoBind and ProMoBind-Base rank second and third overall, outperforming MONN [29], HoTS [30], and BlendNet (T) [31], which also leverage protein-ligand interaction information to improve binding affinity prediction. Notably, ProMoBind-Base, which does not utilize interaction information, achieves a PCC ranking score comparable to BlendNet (T), the best sequence-based baseline. One possible explanation is that the affinity prediction performance of BlendNet may be affected by the accuracy of its pocket extractor, Pseq2Sites. If the predicted binding regions are inaccurate, the downstream affinity prediction module may be negatively affected. Another factor is that ProMoBind-Base uses a protein foundation model pre-trained on large-scale protein data as the protein encoder, which provides informative representations for binding affinity prediction. We also note that PSH-ML [79] achieves the best overall ranking. However, PSH-ML requires experimentally determined complex structures as input. As discussed in the BlendNet study [31], such structures are typically unavailable in virtual screening scenarios. When docking-generated complex structures are used instead, the performance of PSH-ML drops substantially, with a decline of 13 ranking positions. In contrast, all ProMoBind variants are sequence-based models and are therefore not limited by the availability or quality of 3D complex structures. Finally, although ProMoBind achieves a higher overall PCC ranking score than BlendNet (T), we observe that BlendNet (T) performs better than ProMoBind on CSAR2014 and CSARset1 (Supplementary Fig. 2a). This suggests that explicitly modeling pairwise non-covalent interactions can be beneficial, and exploring such interaction modeling strategies could be a promising direction for future work.

We also evaluated the efficiency and practicality of ProMoBind for large-scale virtual screening by measuring both its training time and inference speed. ProMoBind achieves an average training time of 39.8 minutes, which is overall efficient compared to the baseline training speeds listed in Supplementary Table 8. This highlights the effectiveness of ProMoBind’s architecture, enabling it to be fine-tuned efficiently even when incorporating the pre-trained ProMoSite and the foundation models. For inference, ProMoBind processes input sequentially with a batch size of 1 and achieves an average inference time of 0.0012 seconds per protein-ligand pair, making it the fastest model among all baselines, as shown in Supplementary Table 8. This exceptional efficiency underscores ProMoBind’s scalability and practicality for high-throughput virtual screening in real-world drug discovery scenarios.

Since ProMoSite is pre-trained on scPDB, which predominantly consists of protein-ligand complexes involving enzymes, as shown in Supplementary Table 13. To further assess whether the protein family composition of the scPDB pre-training dataset influences generalization across different protein families, we performed an additional protein family–level analysis on the PDBbind v2020 test set under the time-based split [9, 23]. Test samples were grouped by protein family, and only families with more than 10 entries were included to ensure statistical reliability. As shown in Supplementary Table 7, ProMoBind consistently outperforms ProMoBind-Base across all evaluated protein families and evaluation metrics. These results indicate that the performance gains from binding-site–guided pre-training are observed across diverse protein families rather than being confined to a single dominant class, suggesting that our pre-training strategy captures transferable microscale protein–ligand interaction patterns.

### 2.5 ProMoBind enables out-of-distribution binding kinetics prediction

Binding kinetics prediction is an essential yet relatively underexplored aspect of drug discovery and protein–ligand interaction studies, despite growing evidence that drug efficacy depends not only on the strength but also on the speed and duration of drug–target interactions [80–82]. Accurate estimation of binding rates provides key insights into ligand residence time and plays a crucial role in optimizing lead compounds during drug development. We therefore further focus on binding kinetics prediction to evaluate ProMoBind’s performance in downstream tasks of predicting binding characteristics. In our experiment, we collected dissociation rate constant (k_off_) data from PDBbind-koff-2020 [83] and BindingDB [84]. Each sample in the dataset consists of a ligand structure, a protein sequence, and an experimentally determined k_off_ value.

We validated the model’s generalization capability in out-of-distribution (OOD) scenarios using two different data splitting strategies. First, we applied a ligand-based random scaffold split with an 8:1:1 ratio for the training, validation, and test sets, where scaffolds are defined using the Bemis–Murcko method [85], which captures the core structural backbone of small-molecule compounds by retaining the ring systems and linker atoms, ensuring that the test set contains ligands with distinct scaffolds not present in the training or validation sets. For comparison, we adopted the random forest (RF) model and XGBoost [86] as baselines, which are typical methods for binding kinetics prediction [82, 83]. Specifically, the protein inputs were derived from ESM-2 embeddings, and the ligand inputs were represented using either ECFP4 (Extended-Connectivity Fingerprints) [52] or Uni-Mol embeddings. The comparison results, summarized in Table 2 and Supplementary Table 9, show that ProMoBind achieves the best performance across all metrics. Notably, even when only using the embeddings from foundation models, ProMoBind-Base outperforms the RF baseline and achieves performance comparable to XGBoost, demonstrating the effectiveness of our architecture for binding kinetics prediction.

**Table 2.**
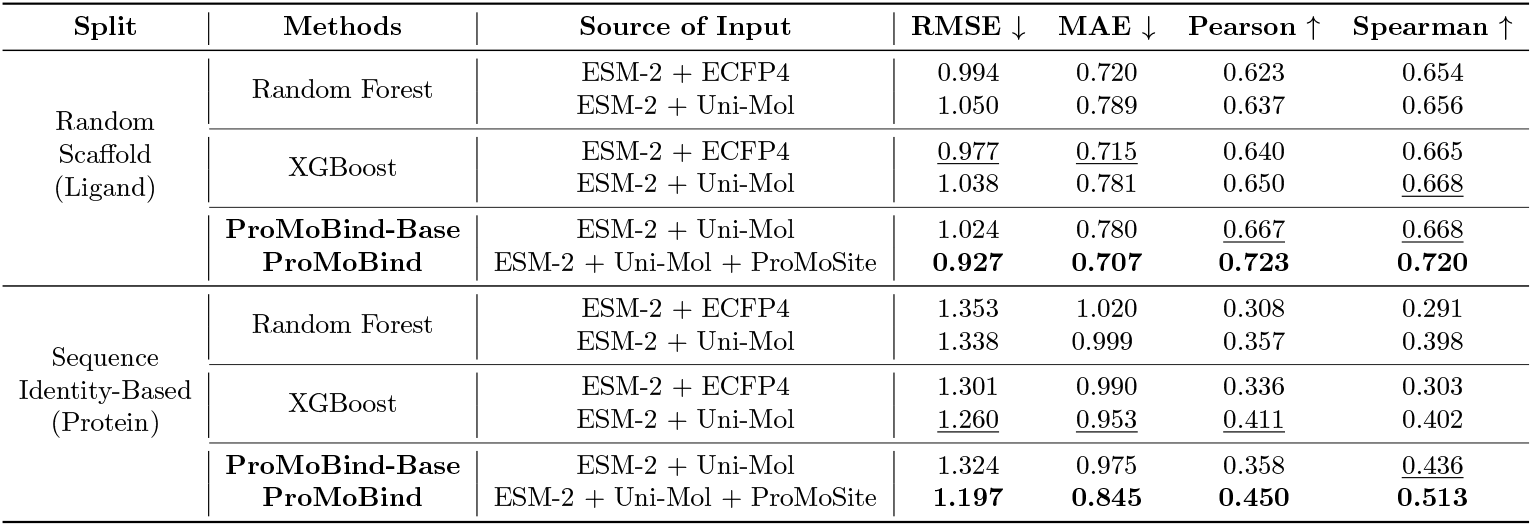
Comparison of model performance in predicting *k*_off_ under different data splits. The source of input for each model is indicated. The best-performing results are highlighted in bold, while the second-best results are underlined.

Second, we evaluated generalization under a protein sequence identity-based split, where we ensured that no protein in the training or validation set shares more than 80% sequence identity with any protein in the test set. As shown in Table 2 and Supplementary Tables 10, ProMoBind again outperforms all baselines across all metrics. Notably, ProMoBind-Base still exceeds the performance of the RF baselines, demonstrating the power of our architecture. Furthermore, the superiority of ProMoBind over ProMoBind-Base demonstrates the effectiveness of the pre-trained ProMoSite on downstream complex-level tasks, and suggests that binding-site-guided pre-training captures interaction patterns that are relevant not only to binding strength but also to dissociation behavior.

These findings highlight ProMoBind’s robust generalization capability and its potential for accurate binding kinetics prediction in OOD settings, further extending its practical value for real-world drug discovery applications.

## 3 Conclusion

In this study, we have presented ProMoNet, a sequence-based pre-training and finetuning framework, addressing key challenges in protein-ligand binding characteristic prediction. We introduce a pre-training strategy based on protein-ligand binding site prediction, which helps connect protein- and ligand-level representations from foundation models to support downstream prediction tasks involving protein-ligand complexes.

ProMoSite, the pre-training module of ProMoNet, excels in identifying both exposed and cryptic binding sites. Its design leverages protein contact maps to model dynamic structural information, avoiding the need for resource-intensive 3D structural modeling or MD simulations. This capability enables ProMoSite to achieve comparable or superior performance to state-of-the-art methods while maintaining scalability and efficiency. Particularly in cryptic binding site prediction, ProMoSite demonstrates its ability to capture dynamic protein-ligand interactions, highlighting the utility of integrating sequence-based features with geometric constraints.

ProMoBind, the fine-tuning module of ProMoNet, incorporates the pre-trained ProMoSite and is optimized for downstream tasks such as binding affinity and kinetics prediction. Its strong performance across diverse benchmarks, including out-of-distribution scenarios, underscores the generalizability and adaptability of this approach. By incorporating ProMoSite’s residue-level binding site predictions, ProMoBind enhances its ability to capture microscale protein-ligand interactions and binding site crypticity, providing an efficient solution for high-throughput compound screening in drug discovery pipelines.

This work underscores the potential of integrating pre-training and fine-tuning strategies in drug discovery. By proposing a solution to bridge protein and molecular foundation models, ProMoNet exemplifies how large-scale datasets can be leveraged to address complex biological tasks. As more high-quality data become available, this approach could be extended to other aspects of protein-ligand interactions, accelerating innovation in computational drug discovery.

While ProMoNet achieves promising results, certain limitations are worth further exploration. Although ProMoNet benefits from the transferred protein and molecular knowledge encoded in ESM-2 (pre-trained on UniRef50) and Uni-Mol (pretrained on ZINC and ChEMBL), our pre-training module, ProMoSite, is trained on scPDB, which predominantly consists of protein-ligand complexes involving enzymes, as shown in Supplementary Table 13. At the same time, some important protein families like GPCRs, ion channels, and membrane receptors remain underrepresented in the datasets used for our evaluation, which limits statistically meaningful, family-specific quantitative analysis in the current study. These protein families often involve large-scale conformational rearrangements, membrane-associated environments, and allosteric regulation, which may require complementary pre-training signals or family-specific adaptation strategies. As future directions, systematic evaluation and potential architectural extensions will be necessary to fully assess its applicability to these important protein families. In addition, our modeling of microscale protein-ligand interactions relies on the accuracy of protein contact maps predicted by ESM-2. For intrinsically disordered proteins, which lack stable tertiary structures, these predictions can be unreliable, potentially limiting the performance of ProMoNet in such cases. Moreover, the reliance on sequence-based inputs, though advantageous for scalability, may limit its ability to capture high-resolution structural details required for certain applications. Future work could investigate the integration of computationally predicted 3D protein structures to complement the current approach. Additionally, extending the framework to the protein-protein binding scenario could further broaden its impact.

In conclusion, ProMoNet provides a pre-training and fine-tuning framework that combines scalability, efficiency, and strong performance. This unified framework offers an effective approach for bridging computational models and practical applications in drug development.

## 4 Methods

### 4.1 ProMoSite architecture

#### 4.1.1 Featurization

For protein, ProMoSite takes protein sequences as input. For proteins with multiple chains, each chain is processed independently together with the ligand as a single-chain protein-ligand pair. Leveraging a pre-trained ESM-2 model (ESM-2-650M) [35] as the protein encoder Encoder_*p*_, the representation of the *i*-th amino acid residue in each provided protein sequence 𝒜^*p*^ = {*a*_*i*_} can be computed: 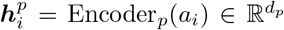, where 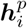 is the *i*-th residue representation, *a*_*i*_ is the amino acid type of the *i*-th residue and *d*_*p*_ is the hidden dimension size. We use the residue-level protein representations from the final layer of ESM-2-650M in all experiments. Additionally, we make use of the unsupervised contact prediction results obtained from ESM-2 to predict residue-residue contact map of each protein 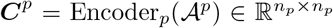, where *n*_*p*_ is the number of residues in 𝒜^*p*^. A contact is defined as a pair of amino acids whose C-*α* distance is less than 8Å.

For ligand, ProMoSite takes SMILES strings [37] as input and uses RDKit [38] to initialize their 3D conformers 𝒢^*l*^. Following the setting in Uni-Mol [36], for each SMILES string, we generate a single random 3D conformer, controlled by a random seed. Then we use the pre-trained Uni-Mol [36] as the ligand encoder Encoder_*l*_ to compute the representation of all non-hydrogen atoms in each ligand conformer: 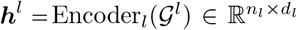, where *d*_*l*_ is the hidden dimension size and *n*_*l*_ is the number of atoms in 𝒢^*l*^. We use 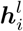 to denote the representation of the *i*-th atom. Besides, the distance map 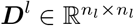 of each 𝒢^*l*^ is computed based on the 3D coordinates of all non-hydrogen atoms in the ligand, where 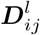 denotes the distance between the *i*-th and *j*-th atom of 𝒢^*l*^ in 3D Euclidean space. Distances greater than 15Å are truncated to 15Å.

We further encode the contact probabilities in ***C***^*p*^ and distances in ***D***^*l*^ using the radial basis function (RBF) in [87] and followed with a multi-layer perception (MLP):

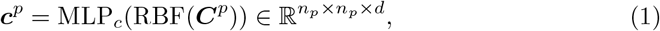

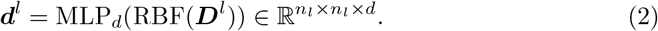

Based on the protein residue representations 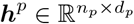 and ligand atom representations 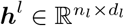, we compute the interaction embedding 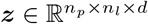 for each protein-ligand pair, which models the interactions between *n*_*p*_ protein residues and *n*_*l*_ ligand atoms. The embedding is a learned numerical representation that encodes relevant features of the input. In ***z***, for ***z***_*ij*_ between the *i*-th protein residue and *j*-th ligand atom, we have:

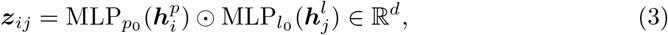

where ⊙ is the element-wise multiplication. Then ***z, c***^*p*^, and ***d***^*l*^ are passed through layer normalization [88] for further updates.

To enable mini-batch training and inference (i.e., processing multiple samples together for computational efficiency) with varying numbers of protein residues and ligand atoms, input embeddings of the same type are zero-padded along their varying dimension to match the maximum length of that dimension within each mini-batch, which is also the approach used by foundation models like ESM-2 [35] and Uni-Mol [36]. For example, consider a mini-batch of *n* protein embeddings, each with hidden dimension *D*, where the maximum number of residues is *L*. After zero-padding, each embedding has a size of *L* × *D* and they are organized into an embedding of size *n* × *L* × *D*, which is then fed into the model. Other inputs, including sets of ligand embeddings {***h***^*l*^}, contact maps {***C***^*p*^}, and distance maps {***D***^*l*^}, are processed in the same way. The padded positions are masked out during subsequent computations, ensuring that only valid entries contribute to the model’s predictions.

#### 4.1.2 Interaction module

Our interaction module (Fig. 1a) contains three steps of updates including *triangle update, triangle self-attention*, and *transition* inspired by [9, 39] to update ***z*** by satisfying the triangle inequality constraints in protein contact maps ***C***^*p*^ and ligand distance maps ***D***^*l*^.

In the *triangle update* step, we update the interaction embedding ***z***_*ij*_ by combining information within each triangle of edges *ij, ik*, and *jk*, where *i* is the *i*-th protein residue, *j* is the *j*-th ligand atom, and *k* can be either protein residue or ligand atom. In each hidden layer of the interaction module, we have:

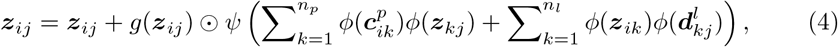

where *g*(·) = sigmoid(Linear(·)), *ψ*(·) = Linear(LayerNorm(·)), and *ϕ*(·) = *g*(·) ⊙ Linear(·). In Equation 4, the first summation term corresponds to the case when *i, k* are protein residues and *j* is a ligand atom, while the second summation term corresponds to the case when *i* is protein residue and *j, k* are ligand atoms.

In the *triangle self-attention* step, we focus on updating the interaction embedding ***z***_*ij*_ between the *i*-th protein residue and the *j*-th ligand atom by considering the effect of all interaction embeddings between the *i*-th protein residue and each ligand atom. By doing so, we can account for the essential physical and chemical effects in protein-ligand interactions, such as steric constraints (excluded-volume, van der Waals interactions) and saturation effects [9]. For example, if the *i*-th protein residue is in close contact with the *j*-th ligand atom, steric constraints limit additional atoms from forming similar close contacts with that residue. Similarly, if the *i*-th residue forms a hydrogen bond with the *j*-th ligand atom, other ligand atoms are less likely to simultaneously form hydrogen bonds with the same residue due to the limited number of hydrogen donors or acceptors. These effects provide a useful framework for modeling microscale protein-ligand interactions.

In each hidden layer, we update ***z***_*ij*_ as follows:

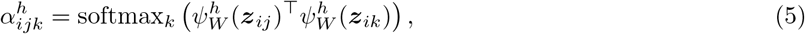

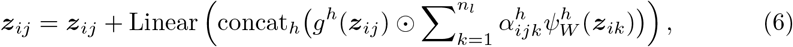

where *h* is the number of attention heads, *ψ*_*W*_ (·) = LinearNoBias(LayerNorm(·)), concat denotes concatenation.

In the *transition* step, we further refine the interaction embedding ***z***_*ij*_ by normalization and nonlinear transformation:

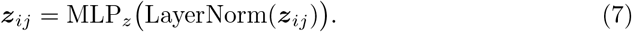

#### 4.1.3 Pooling module

After having the updated interaction embedding 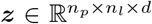, we design a pooling module to compute the score *s* ∈ ℝ for each protein residue, representing the likelihood that the residue belongs to a binding site for the given ligand. We first use a linear transformation to reduce the last dimension size of ***z*** to 1 and then apply sum pooling over the dimension related to ligand atoms to compute a score *s*_1_ for each protein residue. Additionally, another residue-level score, *s*_2_, is computed directly using the protein residue representations ***h***^*p*^ and ligand atom representations ***h***^*l*^. The final score *s* for each residue is obtained by combining *s*_1_ and *s*_2_. For the *i*-th residue, we have:

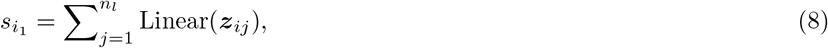

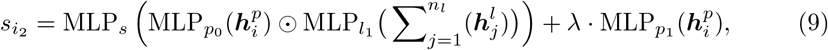

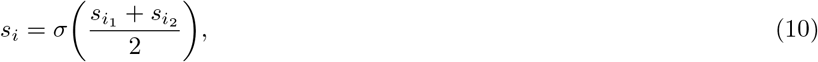

where 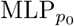 shares the same parameters as the 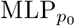 used in the interaction module, *λ* is a scalar value, and *σ*(·) denotes the sigmoid function.

In cases where no ligand is provided as input, such as when predicting cryptic pockets on the PocketMiner dataset [14], which includes some rigid proteins without bound ligands, the score is computed as 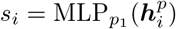.

#### 4.1.4 Clustering and ranking

Although the scores *s* from the pooling module can be directly used for residue-level evaluation (e.g. ROC-AUC and PR-AUC) of ProMoSite, it is also necessary to detect the pockets in 3D space and have pocket-level evaluation (e.g. DCA) to be compared with 3D structure-based methods. To detect 3D pockets, we first normalize all scores 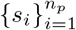 of each protein to the range of 0 to 1. Following this, we apply a threshold *t* to filter out residues that have smaller *s*. With the remaining residues, we utilize the ESM-2 predicted contact map ***C***^*p*^ to define the similarities between each pair of residues and construct a dendrogram that contains the hierarchical clustering information of the residues using the single linkage algorithm [89]. By cutting at a specific linkage distance cutoff *c*, we define the clusters as pocket candidates and then rank them based on the squared sum of the associated residue scores in each pocket candidate. The top-ranked ones are the detected pockets for pocket-level evaluation. For any protein with multiple chains, each chain is processed independently together with the ligand to identify pocket candidates, and the resulting candidates from all chains are then combined and ranked at the protein level. An illustration of these steps can be found in Supplementary Fig. 3.

### 4.2 ProMoBind architecture

#### 4.2.1 Featurization

As shown in Fig. 1b, ProMoBind uses the same way as in ProMoSite to get protein residue representations ***h***^*p*^ and ligand atom representations ***h***^*l*^ from the pre-trained ESM-2 and the pre-trained Uni-Mol, correspondingly. Moreover, ProMoBind takes the predicted scores 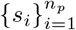 of each protein from the pre-trained ProMoSite as input.

#### 4.2.2 Fusion

To predict the binding characteristic *y* ∈ ℝ between each protein-ligand pair, we first compute the global representation of the protein and ligand, 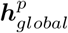 and 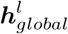, respectively, and then fuse them in the following way:

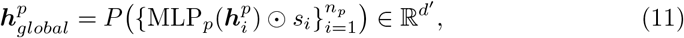

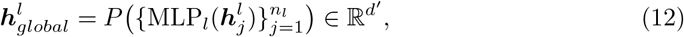

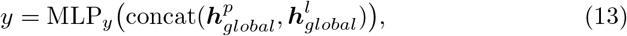

where 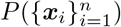 denotes pooling over ***x***_*i*_ for all *i* from 1 to *n*, which can be sum or mean pooling.

Specifically in Equation 11, when pooling all residue representations 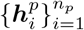 in a protein, we utilize the ProMoSite predicted scores 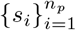 as a gating mechanism to modulate the contribution of each ***h***^*p*^ in 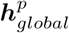 through element-wise multiplication. This approach allows the *s* to selectively emphasize or suppress each ***h***^*p*^ depending on its predicted likelihood to be in a binding site.

### 4.3 Datasets and data splitting strategies

Figure 1d illustrates the experimental setup and dataset usage for evaluating the two modules of ProMoNet, ProMoSite and ProMoBind. In our experiments across different tasks, we apply data splits based on protein sequence identity and/or ligand similarity to avoid data leakage. Supplementary Table 12 summarizes the dataset statistics, data splitting strategies addressing data leakage, and the direct outputs of each module across all experiments in this study. The following sections describe each dataset and the data splitting strategies used in our experiments in detail.

#### 4.3.1 General binding site prediction

Following [44, 47], we use the scPDB v.2017 database [33] and adopt the same data splits for training and cross-validation of ProMoSite. Specifically, the dataset is cleaned and split into 10 cross-validation sets based on their Uniprot IDs to avoid data leakage. We train 10 separate models, each using 9 folds for training and 1 fold for validation. The best set of hyperparameters is selected based on the highest averaged F1 score achieved across the 10 models. This optimal set of hyperparameters is then used to train a final model on the full dataset for evaluation on the test sets. For each protein-ligand pair in scPDB v.2017, the protein is split into single chains, forming pairs between each single chain and each ligand. Irrelevant pairs that are not present in BioLiP [34], a curated database of biologically relevant protein-ligand binding interactions, are removed.

For evaluation, we use COACH420 and HOLO4K [8]. Consistent with [8, 44], we evaluate ProMoSite on the Mlig subsets of these datasets, which contain the relevant ligands for binding sites. The final test sets consist of 291 proteins and 359 ligands for COACH420, and 3413 proteins and 4288 ligands for HOLO4K. To avoid data leakage, we preprocess the training set following the approach in [44] that removes all proteins from the training set that have either sequence identity greater than 50% or ligand similarity greater than 0.9 and sequence identity greater than 30% to any structure in the test set. Additionally, we exclude training pairs where protein sequences exceed 1280 amino acids. After preprocessing, the training set consists of 17101 pairs for COACH420 and 10626 pairs for HOLO4K.

For interaction site prediction, we train ProMoSite on our training set and use MISBD [31] as the test set, which contains 4,778 protein–ligand pairs. Specifically, we exclude the complexes in our scPDB v.2017 training set that overlap with the MISBD test set based on their unique (PDB ID–Ligand ID) pairs, resulting in a training set of 18,082 single chain–ligand pairs. Following our training strategy and hyperparameters in the pre-training task, we use the PocketMiner validation set [14] to select the best ProMoSite model based on the lowest validation loss.

#### 4.3.2 Cryptic binding site prediction

To train ProMoSite, we use the scPDB v.2017 database and preprocess the data to single chain-ligand pairs filtered by BioLiP similar to our preprocessing for general binding site prediction. For validation and testing, we adopt the PocketMiner validation and test sets collected by [14] for cryptic binding site prediction. These sets include proteins with ligand-binding cryptic pockets or rigid structures, comprising 28 and 38 protein-ligand binding pairs or rigid proteins in the validation and test sets, respectively. The best model is selected based on the lowest validation loss. To avoid data leakage, we remove the proteins in the training set that have sequence identity greater than 30% to any of the proteins in the validation and test sets. Additionally, training pairs with protein sequences exceeding 1280 amino acids are removed, resulting in a training set of 19254 single chain-ligand pairs.

For comparison with baselines that predict residue-level probabilities, we use the full PocketMiner test set, which includes both protein-ligand pairs and rigid proteins. For baselines that predict 3D complexes or 3D binding sites, we use a subset of the PocketMiner test set filtered by NeuralPLexer, consisting solely of proteins with ligand-binding cryptic pockets. This subset comprises 19 protein-ligand pairs. Only *apo* protein structures or sequences are used as input.

#### 4.3.3 Pre-training task

In the pre-training task, we pre-train ProMoSite for binding site prediction on the BioLiP-filtered scPDB v.2017 database following a similar process as described in previous ones. The processed training set contains 20403 single chain-ligand pairs. During pre-training, we use the PocketMiner validation set to select the best ProMoSite model based on the lowest validation loss, which is then used to generate scores as inputs for ProMoBind.

#### 4.3.4 Binding affinity prediction

For the experiment on PDBbind v2020 dataset [75], we train and evaluate ProMoBind using two data splits: First, we follow the same time-based split in [9, 23], using data published before 2019 for training and validation, and data published during or after 2019 for testing. For multimers, we concatenate single chain sequences into a single sequence, following the process described in [23]. Sequences exceeding 3,000 amino acids are excluded. The final datasets have 17717, 961, and 363 protein-ligand pairs in the training, validation, and test sets, respectively. Second, we create a protein sequence identity-based split by removing proteins from our time-based training and validation sets that have more than 80% sequence identity with any protein in the test set. The resulting datasets have 15454, 849, and 363 protein-ligand pairs in the training, validation, and test sets, respectively.

For the experiment on the datasets in [31], we train and evaluate ProMoBind using the same datasets and data splits provided in [31] for pair comparison.

#### 4.3.5 Binding kinetics prediction

We curate our own *k*_off_ dataset from PDBbind-koff-2020 [83] and BindingDB [84] for binding kinetics prediction. Inconsistent entries, such as cases where *k*_off_/*k*_on_ values are not consistent with *K*_d_, are removed. After filtering, 632 items from BindingDB and 669 items from PDBbind-koff-2020 are retained. To eliminate redundancy between the two datasets, Tanimoto coefficients (TC) are calculated for the ligands. Redundant pairs, defined as complexes involving the same protein and chemically similar ligands (TC≥0.99), are excluded. Additional filtering removes protein-ligand pairs with mutated protein sequences or multiple *k*_off_ values, resulting in 911 unique pairs. Proteins with sequences longer than 700 amino acids are excluded. After all preprocessing steps, a final dataset of 683 protein-ligand pairs is used for *k*_off_ prediction.

In our experiments, we apply two data splitting strategies for OOD scenarios: First, we use the widely used random scaffold split [90] implemented in RDKit [38] based on the Bemis–Murcko method [85] with an 8:1:1 ratio for the training, validation, and test sets, ensuring that the test set contains ligands with distinct scaffolds not present in the training or validation sets. Second, we employ a protein sequence identity-based split by clustering all proteins in our curated dataset based on their sequence identities to guarantee that each cluster will have less than 80% sequence identity with any other cluster. The largest cluster, which has 93 unique sequences and 520 protein-ligand pairs, is defined as the training set. For the other 27 clusters that have 163 pairs, we randomly split them 5 times at the cluster level by 1:1 to be validation and test sets. As a result, there are no proteins in the training and validation sets that have more than 80% sequence identity with any protein in the test set.

### 4.4 Experimental setting

#### 4.4.1 Implementation Details

In ProMoSite, the hyperparameters of radial basis function (RBF) used in Equation 1 and 2 are identical to those in [87]. The multi-layer perceptrons (MLPs) that encode the results of the RBF function in these equations consist of two linear layers and utilize the self-gated Swish activation function [91]. All other MLPs in ProMoSite and ProMoBind employ ReLU as the activation function and include layer normalization preceding the linear layers. During training, dropout is applied to the triangle update and triangle self-attention steps in ProMoSite to enhance generalization. Further details regarding hyperparameter configurations are provided in Supplementary Table 11.

ProMoSite is trained by minimizing the binary cross-entropy loss between the predicted residue scores and the ground-truth labels. Residues labeled as “unknown” or “unclassified” are excluded from the loss calculation on the PocketMiner dataset. ProMoBind is trained to minimize the mean squared error (MSE) loss between the predicted values and the ground-truth binding characteristics. Both models are optimized using the Adam optimizer.

To enhance memory efficiency while maintaining accuracy, we use PyTorch’s Automatic Mixed Precision (AMP) package for mixed-precision training in ProMoNet. All experiments are conducted on an NVIDIA L40S GPU (48 GB). UCSF ChimeraX [92] is used to generate the visualizations of complexes shown in Fig. 2 and 3.

#### 4.4.2 Baselines

For general binding site prediction, we compare ProMoSite with baseline results reported in [44] for fair comparison. We retrain Pseq2Sites using our training set and training strategy based on the code at https://github.com/Blue1993/Pseq2Sites. Additionally, we evaluate PUResNet, DeepProSite, and HoTS using their publicly available trained models. PUResNet is accessible at https://github.com/jivankandel/PUResNet, DeepProSite at https://github.com/WeiLab-Biology/DeepProSite, and HoTS at https://github.com/GIST-CSBL/HoTS. For interaction site prediction, we adopt the baseline results reported in [31].

For cryptic binding site prediction, we use the results of CryptoSite and PocketMiner reported in [14]. Other baseline models, including NeuralPLexer-AF2, NeuralPLexer-fasta, DynamicBind, and P2Rank, are evaluated using their publicly available trained models. NeuralPLexer variants are available at https://github.com/zrqiao/NeuralPLexer, DynamicBind at https://github.com/luwei0917/DynamicBind, and P2Rank (version 2.4.2) at https://github.com/rdk/p2rank. When measuring inference speed, input data is processed sequentially for each model to ensure fair comparison, and preprocessing time is excluded.

For binding affinity prediction, we adopt the baseline results reported in [17, 23, 67] under the time-based split on PDBbind v2020. The results of Boltz-2 are obtained by performing inference on our test set with its publicly available pretrained model at https://github.com/jwohlwend/boltz. For the sequence identity-based split, we obtain the results of PSICHIC by training the model using the publicly available code (https://github.com/huankoh/PSICHIC) and the hyperparameters provided by the original authors. For the comparison with the baselines on the eight test datasets in [31], we adopt the baseline results reported in [31].

For binding kinetics prediction, in all experiments, we implement the random forest baselines using the RandomForestRegressor from the scikit-learn library. We perform hyperparameter optimization through grid search. The optimized hyperparameters include max depth, n estimators, min samples split, and min samples leaf. For XGBoost, we employ the XGBRegressor from the xgboost library. A grid search is conducted to optimize the hyperparameters, including n estimators, max depth, learning rate, min child weight, subsample, colsample bytree, reg alpha, reg lambda, and gamma, based on validation performance. ECFP4 fingerprints are computed with RDKit using a 1024-bit setting.

### 4.5 Relationship to Prior Methods

Several previous studies have explored incorporating protein–ligand interaction information to improve binding affinity prediction, including MONN [29], HoTS [30], and BlendNet [31]. Although these studies share the general idea of leveraging interaction signals, the architectural design, implementation strategies, and scope of ProMoNet differ substantially from those of prior approaches, and ProMoNet also addresses several limitations of these methods. A comparison between these methods and ProMoNet is provided in Supplementary Table 2 and discussed in detail below.

#### Model architectures

Our encoders are built upon foundation models for both proteins and ligands, providing substantially broader data coverage and more diverse representations. In contrast, prior models largely rely on task-specific encoders trained from scratch, with BlendNet being the only method among them that incorporates a ligand encoder pretrained on large-scale molecular data.

#### Strategies to model protein–ligand interactions

MONN and HoTS model interactions between the ligand and the full protein sequence. BlendNet instead relies on an external binding pocket prediction tool (Pseq2Sites) to first identify binding pockets and then models interactions between ligand and predicted pocket subsequences. Consequently, the performance of BlendNet is closely tied to the accuracy of the pocket prediction. If the predicted pockets do not cover the true binding region, the subsequent interaction modeling and affinity prediction may be negatively affected. As shown in Fig. 2b, Pseq2Sites achieves lower performance compared with our proposed ProMoSite. Moreover, unlike ProMoSite, which predicts ligand-specific binding sites, Pseq2Sites predicts binding pockets solely from the protein sequence without considering ligand information. In practice, different ligands may bind to different sites on the same protein, which may limit the effectiveness of BlendNet in virtual screening when Pseq2Sites is used to identify protein pockets.

In addition, HoTS splits each protein sequence into grids (binding regions) and models interactions between ligands and these regions, which lead to relatively coarse-grained binding representations. In contrast, ProMoSite models interactions between each ligand atom and each protein residue, enabling a more fine-grained representation of protein–ligand binding patterns. Moreover, the interaction module in ProMoSite incorporates geometric constraints derived from the triangle inequality to better preserve spatial consistency in the learned representations. Such geometric constraints help the model capture underlying physical and chemical relationships between residues and ligand atoms. MONN, HoTS, and BlendNet do not explicitly incorporate this type of geometric constraint in their interaction modeling frameworks.

#### Approaches to utilize interaction information for binding affinity prediction

MONN, HoTS, and BlendNet all require the joint supervision of protein–ligand interaction labels (such as pairwise non-covalent interactions or binding regions) and binding affinity labels during training. In contrast, our framework introduces a pretraining stage that uses only binding site labels to train ProMoSite. In downstream tasks such as binding affinity prediction with ProMoBind, the pretrained ProMoSite is kept frozen, and the predicted binding site scores are incorporated as inputs to ProMoBind to transfer the learned knowledge. This design allows ProMoBind to be trained efficiently and flexibly using only binding affinity labels.

Furthermore, MONN and BlendNet (T) require training datasets in which protein–ligand pairs are annotated with both binding affinity labels and pairwise non-covalent interaction labels derived from experimentally determined complex structures. However, many widely used binding affinity datasets (e.g., ChEMBL [93]) do not provide such structures, which makes it difficult to derive reliable pairwise interaction labels. As a result, these models are restricted to relatively limited datasets. In contrast, ProMoBind can be trained directly on datasets that only contain binding affinity labels, allowing it to utilize a broader range of available data.

#### Scopes of tasks

MONN and BlendNet focus on predicting pairwise non-covalent interactions and binding affinity, while HoTS performs coarse-grained binding region prediction and binding affinity prediction. In comparison, our proposed ProMoNet framework can detect general binding sites and predict binding affinity, as well as cryptic binding site detection and binding kinetics prediction, which are not covered in the aforementioned works.

## Declarations

### Availability of data and materials

The datasets used in this study are publicly available. scPDB v.2017 database is accessible at http://bioinfo-pharma.u-strasbg.fr/scPDB/, and BioLiP database can be found at https://zhanggroup.org/BioLiP/. Mlig subsets of COACH420 and HOLO4K, as well as indices for the 10-fold cross-validation split of scPDB, are available at https://github.com/devalab/DeepPocket. MISBD dataset is available at https://github.com/Blue1993/BlendNet. Pocket-Miner validation and test sets are accessible via https://www.nature.com/articles/s41467-023-36699-3#Sec34 and https://github.com/Mickdub/gvp/tree/pocket_pred/data/pm-dataset. *Apo*-*holo* pair systems from PocketMiner dataset used by NeuralPLexer are available at https://zenodo.org/records/10373581. PDBbind v2020 dataset and its time-based split are accessible at https://github.com/huankoh/PSICHIC/tree/main/dataset/pdb2020. The training set and the eight test datasets used for binding affinity prediction in [31] are available at https://github.com/Blue1993/BlendNet. PDBbind-koff-2020 database can be downloaded from http://www.pdbbind.org.cn/download/koff_dataset.rar, and BindingDB database is publicly accessible at https://www.bindingdb.org/. The source code of this work is available at https://github.com/zetayue/ProMoNet.

### Authors’ contributions

S. Z. conceived the concept, designed the method and the experiments, prepared data, implemented the algorithms, performed the experiments, analyzed results, and drafted the manuscript. Li X. prepared data and performed the baseline experiments for binding kinetics prediction. D. T. performed the baseline experiments for binding site prediction. Li X. and Lei X. provided edits to the manuscript. Lei X. conceived the concept, designed the method and the experiments, and acquired funding.

## Acknowledgements

We would like to thank Amitesh Badkul for contributions to the binding affinity prediction task, including insightful discussions and experimental support that helped refine our approach.

## References

[1] Kitchen, D.B., Decornez, H., Furr, J.R., Bajorath, J.: Docking and scoring in virtual screening for drug discovery: methods and applications. Nature reviews Drug discovery 3(11), 935–949 (2004)

[2] Śledź, P., Caflisch, A.: Protein structure-based drug design: from docking to molecular dynamics. Current opinion in structural biology 48, 93–102 (2018)

[3] Perozzo, R., Folkers, G., Scapozza, L.: Thermodynamics of protein–ligand interactions: history, presence, and future aspects. Journal of Receptors and Signal Transduction 24(1-2), 1–52 (2004)

[4] Copeland, R.A., Pompliano, D.L., Meek, T.D.: Drug–target residence time and its implications for lead optimization. Nature reviews Drug discovery 5(9), 730–739 (2006)

[5] Du, X., Li, Y., Xia, Y.-L., Ai, S.-M., Liang, J., Sang, P., Ji, X.-L., Liu, S.-Q.: Insights into protein–ligand interactions: mechanisms, models, and methods. International journal of molecular sciences 17(2), 144 (2016)

[6] Dhakal, A., McKay, C., Tanner, J.J., Cheng, J.: Artificial intelligence in the prediction of protein–ligand interactions: recent advances and future directions. Briefings in Bioinformatics 23(1), 476 (2022)

[7] Cai, C., Wang, S., Xu, Y., Zhang, W., Tang, K., Ouyang, Q., Lai, L., Pei, J.: Transfer learning for drug discovery. Journal of Medicinal Chemistry 63(16), 8683–8694 (2020)

[8] Krivák, R., Hoksza, D.: P2rank: machine learning based tool for rapid and accurate prediction of ligand binding sites from protein structure. Journal of cheminformatics 10, 1–12 (2018)

[9] Lu, W., Wu, Q., Zhang, J., Rao, J., Li, C., Zheng, S.: Tankbind: Trigonometry-aware neural networks for drug-protein binding structure prediction. Advances in neural information processing systems 35, 7236–7249 (2022)

[10] Scardino, V., Di Filippo, J.I., Cavasotto, C.N.: How good are alphafold models for docking-based virtual screening? Iscience 26(1) (2023)

[11] Wang, D.D., Ou-Yang, L., Xie, H., Zhu, M., Yan, H.: Predicting the impacts of mutations on protein-ligand binding affinity based on molecular dynamics simulations and machine learning methods. Computational and structural biotechnology journal 18, 439–454 (2020)

[12] Decherchi, S., Cavalli, A.: Thermodynamics and kinetics of drug-target binding by molecular simulation. Chemical Reviews 120(23), 12788–12833 (2020)

[13] Kuzmanic, A., Bowman, G.R., Juarez-Jimenez, J., Michel, J., Gervasio, F.L.: Investigating cryptic binding sites by molecular dynamics simulations. Accounts of chemical research 53(3), 654–661 (2020)

[14] Meller, A., Ward, M., Borowsky, J., Kshirsagar, M., Lotthammer, J.M., Oviedo, F., Ferres, J.L., Bowman, G.R.: Predicting locations of cryptic pockets from single protein structures using the pocketminer graph neural network. Nature Communications 14(1), 1177 (2023)

[15] Wang, J., Do, H.N., Koirala, K., Miao, Y.: Predicting biomolecular binding kinetics: A review. Journal of Chemical Theory and Computation 19(8), 2135–2148 (2023)

[16] Qiao, Z., Nie, W., Vahdat, A., al.: State-specific protein–ligand complex structure prediction with a multiscale deep generative model. Nature Machine Intelligence, 1–14 (2024)

[17] Lu, W., Zhang, J., Huang, W., al.: Dynamicbind: predicting ligand-specific protein-ligand complex structure with a deep equivariant generative model. Nature Communications 15(1), 1071 (2024)

[18] Öztürk, H., Ö zgür, A., Ozkirimli, E.: Deepdta: deep drug–target binding affinity prediction. Bioinformatics 34(17), 821–829 (2018)

[19] Nguyen, T., Le, H., Quinn, T.P., Nguyen, T., Le, T.D., Venkatesh, S.: Graphdta: predicting drug–target binding affinity with graph neural networks. Bioinformatics 37(8), 1140–1147 (2021)

[20] Badkul, A., Xie, L., Zhang, S., Xie, L.: Multimodal out-of-distribution individual uncertainty quantification enhances binding affinity prediction for polypharmacology. Nature Machine Intelligence, 1–11 (2025)

[21] Li, S., Zhou, J., Xu, T., Huang, L., Wang, F., Xiong, H., Huang, W., Dou, D., Xiong, H.: Structure-aware interactive graph neural networks for the prediction of protein-ligand binding affinity. In: Proceedings of the 27th ACM SIGKDD Conference on Knowledge Discovery & Data Mining, pp. 975–985 (2021)

[22] Zhang, S., Liu, Y., Xie, L.: A universal framework for accurate and efficient geometric deep learning of molecular systems. Scientific Reports 13(1), 19171 (2023)

[23] Koh, H.Y., Nguyen, A.T., Pan, S., May, L.T., Webb, G.I.: Physicochemical graph neural network for learning protein–ligand interaction fingerprints from sequence data. Nature Machine Intelligence, 1–15 (2024)

[24] Bommasani, R., Hudson, D.A., Adeli, E., Altman, R., Arora, S., Arx, S., Bernstein, M.S., Bohg, J., Bosselut, A., Brunskill, E., et al.: On the opportunities and risks of foundation models. arXiv preprint 2108.07258 (2021)

[25] Zhang, Q., Ding, K., Lyv, T., Wang, X., Yin, Q., Zhang, Y., Yu, J., Wang, Y., Li, X., Xiang, Z., et al.: Scientific large language models: A survey on biological & chemical domains. arXiv preprint 2401.14656 (2024)

[26] Li, Q., Hu, Z., Wang, Y., Li, L., Fan, Y., King, I., Song, L., Li, Y.: Progress and opportunities of foundation models in bioinformatics. arXiv preprint 2402.04286 (2024)

[27] Wang, X., Chen, G., Qian, G., Gao, P., Wei, X.-Y., Wang, Y., Tian, Y., Gao, W.: Large-scale multi-modal pre-trained models: A comprehensive survey. Machine Intelligence Research 20(4), 447–482 (2023)

[28] Sledzieski, S., Kshirsagar, M., Baek, M., Dodhia, R., Lavista Ferres, J., Berger, B.: Democratizing protein language models with parameter-efficient fine-tuning. Proceedings of the National Academy of Sciences 121(26), 2405840121 (2024)

[29] Li, S., Wan, F., Shu, H., Jiang, T., Zhao, D., Zeng, J.: Monn: a multi-objective neural network for predicting compound-protein interactions and affinities. Cell systems 10(4), 308–322 (2020)

[30] Lee, I., Nam, H.: Sequence-based prediction of protein binding regions and drug– target interactions. Journal of cheminformatics 14(1), 5 (2022)

[31] Seo, S., Kim, H., Lee, J., Choi, S., Park, S.: Exploring the potential of compound– protein complex structure-free models in virtual screening using blendnet. Briefings in Bioinformatics 26(1) (2024)

[32] Vajda, S., Beglov, D., Wakefield, A.E., Egbert, M., Whitty, A.: Cryptic binding sites on proteins: definition, detection, and druggability. Current opinion in chemical biology 44, 1–8 (2018)

[33] Desaphy, J., Bret, G., Rognan, D., Kellenberger, E.: sc-pdb: a 3d-database of ligandable binding sites—10 years on. Nucleic acids research 43(D1), 399–404 (2015)

[34] Yang, J., Roy, A., Zhang, Y.: Biolip: a semi-manually curated database for biologically relevant ligand–protein interactions. Nucleic acids research 41(D1), 1096–1103 (2012)

[35] Lin, Z., Akin, H., Rao, R., Hie, B., Zhu, Z., Lu, W., Smetanin, N., Verkuil, R., Kabeli, O., Shmueli, Y., et al.: Evolutionary-scale prediction of atomic-level protein structure with a language model. Science 379(6637), 1123–1130 (2023)

[36] Zhou, G., Gao, Z., Ding, Q., al.: Uni-mol: A universal 3d molecular representation learning framework. In: The Eleventh International Conference on Learning Representations (2022)

[37] Weininger, D.: Smiles, a chemical language and information system. 1. introduction to methodology and encoding rules. Journal of chemical information and computer sciences 28(1), 31–36 (1988)

[38] Landrum, G.: RDKit: A software suite for cheminformatics, computational chemistry, and predictive modeling. Academic Press (2013)

[39] Jumper, J., Evans, R., Pritzel, A., Green, T., Figurnov, M., Ronneberger, O., Tunyasuvunakool, K., Bates, R., Žídek, A., Potapenko, A., et al.: Highly accurate protein structure prediction with alphafold. Nature 596(7873), 583–589 (2021)

[40] Vassura, M., Margara, L., Di Lena, P., Medri, F., Fariselli, P., Casadio, R.: Reconstruction of 3d structures from protein contact maps. IEEE/ACM Transactions on Computational Biology and Bioinformatics 5(3), 357–367 (2008)

[41] Li, J., Wang, L., Zhu, Z., Song, C.: Exploring the alternative conformation of a known protein structure based on contact map prediction. Journal of Chemical Information and Modeling 64(1), 301–315 (2023)

[42] Vaswani, A., Shazeer, N., Parmar, N., Uszkoreit, J., Jones, L., Gomez, A.N., Kaiser, L., Polosukhin, I.: Attention is all you need. Advances in neural information processing systems 30 (2017)

[43] Su, D., Xu, Y., Winata, G.I., Xu, P., Kim, H., Liu, Z., Fung, P.: Generalizing question answering system with pre-trained language model fine-tuning. In: Proceedings of the 2nd Workshop on Machine Reading for Question Answering, pp. 203–211 (2019)

[44] Aggarwal, R., Gupta, A., Chelur, V., Jawahar, C., Priyakumar, U.D.: Deeppocket: ligand binding site detection and segmentation using 3d convolutional neural networks. Journal of Chemical Information and Modeling 62(21), 5069–5079 (2021)

[45] Le Guilloux, V., Schmidtke, P., Tuffery, P.: Fpocket: an open source platform for ligand pocket detection. BMC bioinformatics 10(1), 1–11 (2009)

[46] Jiménez, J., Doerr, S., Martínez-Rosell, G., Rose, A.S., De Fabritiis, G.: Deepsite: protein-binding site predictor using 3d-convolutional neural networks. Bioinformatics 33(19), 3036–3042 (2017)

[47] Stepniewska-Dziubinska, M.M., Zielenkiewicz, P., Siedlecki, P.: Improving detection of protein-ligand binding sites with 3d segmentation. Scientific reports 10(1), 5035 (2020)

[48] Kandel, J., Tayara, H., Chong, K.T.: Puresnet: prediction of protein-ligand binding sites using deep residual neural network. Journal of cheminformatics 13(1), 1–14 (2021)

[49] Fang, Y., Jiang, Y., Wei, L., Ma, Q., Ren, Z., Yuan, Q., Wei, D.-Q.: Deepprosite: structure-aware protein binding site prediction using esmfold and pretrained language model. Bioinformatics 39(12), 718 (2023)

[50] Seo, S., Choi, J., Choi, S., Lee, J., Park, C., Park, S.: Pseq2sites: enhancing protein sequence-based ligand binding-site prediction accuracy via the deep convolutional network and attention mechanism. Engineering Applications of Artificial Intelligence 127, 107257 (2024)

[51] Redmon, J.: You only look once: Unified, real-time object detection. In: Proceedings of the IEEE Conference on Computer Vision and Pattern Recognition (2016)

[52] Rogers, D., Hahn, M.: Extended-connectivity fingerprints. Journal of chemical information and modeling 50(5), 742–754 (2010)

[53] He, X., Zhao, L., Tian, Y., Li, R., Chu, Q., Gu, Z., Zheng, M., Wang, Y., Li, S., Jiang, H., et al.: Highly accurate carbohydrate-binding site prediction with deepglycansite. Nature Communications 15(1), 5163 (2024)

[54] Pandey, M., Radaeva, M., Mslati, H., Garland, O., Fernandez, M., Ester, M., Cherkasov, A.: Ligand binding prediction using protein structure graphs and residual graph attention networks. Molecules 27(16), 5114 (2022)

[55] Yuan, W., Chen, G., Chen, C.Y.-C.: Fusiondta: attention-based feature polymerizer and knowledge distillation for drug-target binding affinity prediction. Briefings in Bioinformatics 23(1), 506 (2022)

[56] Jin, Z., Wu, T., Chen, T., Pan, D., Wang, X., Xie, J., Quan, L., Lyu, Q.: Capla: improved prediction of protein–ligand binding affinity by a deep learning approach based on a cross-attention mechanism. Bioinformatics 39(2), 049 (2023)

[57] Lu, R., Wang, J., Li, P., Li, Y., Tan, S., Pan, Y., Liu, H., Gao, P., Xie, G., Yao, X.: Improving drug-target affinity prediction via feature fusion and knowledge distillation. Briefings in Bioinformatics 24(3), 145 (2023)

[58] Team, I.L.: Accurate Predictions of Novel Biomolecular Interactions with IsoDDE. (2026). https://doi.org/10.5281/zenodo.18606681. 10.5281/zenodo.18606681

[59] Cimermancic, P., Weinkam, P., Rettenmaier, T.J., al: Cryptosite: expanding the druggable proteome by characterization and prediction of cryptic binding sites. Journal of molecular biology 428(4), 709–719 (2016)

[60] Song, Y., Sohl-Dickstein, J., Kingma, D.P., Kumar, A., Ermon, S., Poole, B.: Score-based generative modeling through stochastic differential equations. arXiv preprint 2011.13456 (2020)

[61] Stepniewska-Dziubinska, M.M., Zielenkiewicz, P., Siedlecki, P.: Development and evaluation of a deep learning model for protein–ligand binding affinity prediction. Bioinformatics 34(21), 3666–3674 (2018)

[62] Zheng, L., Fan, J., Mu, Y.: Onionnet: a multiple-layer intermolecular-contact-based convolutional neural network for protein–ligand binding affinity prediction. ACS omega 4(14), 15956–15965 (2019)

[63] Jiang, D., Hsieh, C.-Y., Wu, Z., Kang, Y., Wang, J., Wang, E., Liao, B., Shen, C., Xu, L., Wu, J., et al.: Interactiongraphnet: A novel and efficient deep graph representation learning framework for accurate protein–ligand interaction predictions. Journal of medicinal chemistry 64(24), 18209–18232 (2021)

[64] Koes, D.R., Baumgartner, M.P., Camacho, C.J.: Lessons learned in empirical scoring with smina from the csar 2011 benchmarking exercise. Journal of chemical information and modeling 53(8), 1893–1904 (2013)

[65] McNutt, A.T., Francoeur, P., Aggarwal, R., Masuda, T., Meli, R., Ragoza, M., Sunseri, J., Koes, D.R.: Gnina 1.0: molecular docking with deep learning. Journal of cheminformatics 13(1), 43 (2021)

[66] Sverrisson, F., Feydy, J., Correia, B.E., Bronstein, M.M.: Fast end-to-end learning on protein surfaces. In: Proceedings of the IEEE/CVF Conference on Computer Vision and Pattern Recognition, pp. 15272–15281 (2021)

[67] Lai, H., Wang, L., Qian, R., Huang, J., Zhou, P., Ye, G., Wu, F., Wu, F., Zeng, X., Liu, W.: Author correction: Interformer: an interaction-aware model for protein-ligand docking and affinity prediction. nature communications 16, 1566 (2025)

[68] Chen, L., Tan, X., Wang, D., Zhong, F., Liu, X., Yang, T., Luo, X., Chen, K., Jiang, H., Zheng, M.: Transformercpi: improving compound–protein interaction prediction by sequence-based deep learning with self-attention mechanism and label reversal experiments. Bioinformatics 36(16), 4406–4414 (2020)

[69] Huang, K., Xiao, C., Glass, L.M., Sun, J.: Moltrans: molecular interaction transformer for drug–target interaction prediction. Bioinformatics 37(6), 830–836 (2021)

[70] Bai, P., Miljković, F., John, B., Lu, H.: Interpretable bilinear attention network with domain adaptation improves drug–target prediction. Nature Machine Intelligence 5(2), 126–136 (2023)

[71] Jiang, M., Li, Z., Zhang, S., Wang, S., Wang, X., Yuan, Q., Wei, Z.: Drug–target affinity prediction using graph neural network and contact maps. RSC advances 10(35), 20701–20712 (2020)

[72] Jiang, M., Wang, S., Zhang, S., Zhou, W., Zhang, Y., Li, Z.: Sequence-based drugtarget affinity prediction using weighted graph neural networks. BMC genomics 23(1), 449 (2022)

[73] Wang, P., Zheng, S., Jiang, Y., Li, C., Liu, J., Wen, C., Patronov, A., Qian, D., Chen, H., Yang, Y.: Structure-aware multimodal deep learning for drug–protein interaction prediction. Journal of chemical information and modeling 62(5), 1308– 1317 (2022)

[74] Passaro, S., Corso, G., Wohlwend, J., Reveiz, M., Thaler, S., Somnath, V.R., Getz, N., Portnoi, T., Roy, J., Stark, H., et al.: Boltz-2: Towards accurate and efficient binding affinity prediction. BioRxiv (2025)

[75] Wang, R., Fang, X., Lu, Y., Yang, C.-Y., Wang, S.: The pdbbind database: methodologies and updates. Journal of medicinal chemistry 48(12), 4111–4119 (2005)

[76] Su, M., Yang, Q., Du, Y., Feng, G., Liu, Z., Li, Y., Wang, R.: Comparative assessment of scoring functions: the casf-2016 update. Journal of chemical information and modeling 59(2), 895–913 (2018)

[77] Carlson, H.A., Smith, R.D., Damm-Ganamet, K.L., Stuckey, J.A., Ahmed, A., Convery, M.A., Somers, D.O., Kranz, M., Elkins, P.A., Cui, G., et al.: Csar 2014: a benchmark exercise using unpublished data from pharma. Journal of chemical information and modeling 56(6), 1063–1077 (2016)

[78] Hartshorn, M.J., Verdonk, M.L., Chessari, G., Brewerton, S.C., Mooij, W.T., Mortenson, P.N., Murray, C.W.: Diverse, high-quality test set for the validation of proteinligand docking performance. Journal of medicinal chemistry 50(4), 726–741 (2007)

[79] Liu, X., Feng, H., Wu, J., Xia, K.: Persistent spectral hypergraph based machine learning (psh-ml) for protein-ligand binding affinity prediction. Briefings in bioinformatics 22(5), 127 (2021)

[80] Copeland, R.A.: Conformational adaptation in drug–target interactions and residence time. Future medicinal chemistry 3(12), 1491–1501 (2011)

[81] Pan, A.C., Borhani, D.W., Dror, R.O., Shaw, D.E.: Molecular determinants of drug–receptor binding kinetics. Drug discovery today 18(13-14), 667–673 (2013)

[82] Amangeldiuly, N., Karlov, D., Fedorov, M.V.: Baseline model for predicting protein–ligand unbinding kinetics through machine learning. Journal of Chemical Information and Modeling 60(12), 5946–5956 (2020)

[83] Liu, H., Su, M., Lin, H.-X., Wang, R., Li, Y.: Public data set of protein–ligand dissociation kinetic constants for quantitative structure–kinetics relationship studies. ACS omega 7(22), 18985–18996 (2022)

[84] Liu, T., Hwang, L., Burley, S.K., Nitsche, C.I., Southan, C., Walters, W.P., Gilson, M.K.: Bindingdb in 2024: a fair knowledgebase of protein-small molecule binding data. Nucleic Acids Research, 1075 (2024)

[85] Bemis, G.W., Murcko, M.A.: The properties of known drugs. 1. molecular frameworks. Journal of medicinal chemistry 39(15), 2887–2893 (1996)

[86] Chen, T., Guestrin, C.: Xgboost. In: Proceedings of the 22nd ACM SIGKDD International Conference on Knowledge Discovery and Data Mining, vol. 13, pp. 785–794 (2016). ACM

[87] Gasteiger, J., Groß, J., Günnemann, S.: Directional message passing for molecular graphs. In: International Conference on Learning Representations (ICLR) (2020)

[88] Lei Ba, J., Kiros, J.R., Hinton, G.E.: Layer normalization. ArXiv e-prints, 1607 (2016)

[89] Müllner, D.: Modern hierarchical, agglomerative clustering algorithms. arXiv preprint 1109.2378 (2011)

[90] Wu, Z., Ramsundar, B., Feinberg, E.N., Gomes, J., Geniesse, C., Pappu, A.S., Leswing, K., Pande, V.: Moleculenet: a benchmark for molecular machine learning. Chemical science 9(2), 513–530 (2018)

[91] Ramachandran, P., Zoph, B., Le, Q.V.: Searching for activation functions. arXiv preprint 1710.05941 (2017)

[92] Meng, E.C., Goddard, T.D., Pettersen, E.F., Couch, G.S., Pearson, Z.J., Morris, J.H., Ferrin, T.E.: Ucsf chimerax: Tools for structure building and analysis. Protein Science 32(11), 4792 (2023)

[93] Mendez, D., Gaulton, A., Bento, A.P., Chambers, J., De Veij, M., Félix, E., Magariños, M.P., Mosquera, J.F., Mutowo, P., Nowotka, M., et al.: Chembl: towards direct deposition of bioassay data. Nucleic acids research 47(D1), 930–940 (2019)

