## Supplementary Information for "Sequence-based Drug-Target Binding Site Pre-training Enables Cryptic Pocket Detection and Improves Binding Affinity and Kinetics Prediction"

#### Supplementary Tables

**Supplementary Table 1** Results of the ablation study on ProMoSite, presented as Top-n Success Rate (SR). In the ablation "w/o Interaction Module", the interaction embedding $\boldsymbol{z}$ is excluded, and consequently, $s_1$ (Eq. 8) is omitted when computing the final score $s$.

| Ablation | COACH420 | HOLO4K |
|---|---|---|
| ProMoSite | **68.30±0.99** | **73.46±0.65** |
| w/o Interaction Module | 66.10±0.92 | 71.33±0.55 |
| w/o $s_2$ (Eq. 9) | 65.58±1.08 | 71.19±0.73 |

**Supplementary Table 2** Model input, architectural components, training strategies, and predictive tasks of related models.

| Model | Input | Ligand Encoder | Protein Encoder | Protein-Ligand Interaction Modeling | Training Strategy | Tasks |
|---|---|---|---|---|---|---|
| MONN | Ligand: molecular graph<br>Protein: full protein sequence | Graph Convolution | CNN | Inner product with sigmoid function | Train the model end-to-end to predict pairwise noncovalent interactions and binding affinity | 1. Pairwise interaction prediction<br>2. Binding affinity prediction |
| HoTS | Ligand: molecular graph<br>Protein: full protein sequence | Morgan/circular fingerprints with fully connected layers | CNN + grid pooling | Transformer blocks | 1. Pre-train the model to predict binding regions<br>2. Fine-tune it to predict binding regions and drug–target interactions | 1. Binding region prediction<br>2. Drug–target interaction prediction |
| BlendNet | Ligand: molecular graph<br>Protein: pocket protein sequence | Pre-trained graph neural network | Multi-head self-attention layer | Multi-head cross-attention | 1. Train a teacher model to predict binding affinities and noncovalent interactions<br>2. Train a student model in two stages under teacher guidance to predict the same two objectives | 1. Noncovalent interaction prediction<br>2. Binding affinity prediction |
| ProMoNet (Ours) | Ligand: SMILES string<br>Protein: full protein sequence | Pre-trained molecular foundation model | Pre-trained protein foundation model | 1. ProMoSite: Triangle update and multi-head cross-attention constrained by triangle inequality<br>2. ProMoBind: Gated by ProMoSite-predicted scores, followed by feature concatenation and MLP | 1. Train ProMoSite to predict exposed and cryptic binding sites<br>2. Train ProMoBind using ProMoSite outputs to predict binding affinity or kinetics | 1. General binding site prediction<br>2. Cryptic binding site prediction<br>3. Binding affinity prediction<br>4. Binding kinetics prediction |

**Supplementary Table 3** Detailed DCA values on PocketMiner test set. DCA values $\geq$ 4Å are highlighted in bold, indicating failed predictions.

| PDB ID | Ligand Code | NeuralPLexer-fasta | NeuralPLexer-AF2 | DynamicBind | P2Rank | ProMoSite |
|---|---|---|---|---|---|---|
| 1KMO | CIT | **6.6179** | 2.2680 | 3.3691 | **31.7803** | **8.0241** |
| 1KX9 | BDD | 2.1874 | 1.4772 | 1.2245 | 2.0727 | 2.3557 |
| 1TVQ | CHD | 1.0457 | 2.3125 | 1.2632 | 1.9921 | 1.0543 |
| 2FJY | PRZ | 2.6492 | 0.9086 | **4.0136** | **4.4056** | 1.5673 |
| 2HQ8 | CTZ | 3.9494 | 3.6171 | 3.5262 | 2.2861 | 3.6782 |
| 2ZKU | JT1 | **4.4851** | **4.9895** | **13.4771** | **7.2699** | **8.4981** |
| 3NX1 | FER | 3.1940 | 1.9785 | **4.5397** | **5.4812** | 3.6931 |
| 3P53 | H0H | **26.0488** | **27.7842** | **16.7689** | **25.1612** | **21.3088** |
| 3UGK | NAI | 3.8310 | 0.6547 | 3.2013 | 3.6811 | 1.3606 |
| 3UGK | SHR | 1.7931 | **6.2229** | 1.6863 | 1.4801 | **5.0005** |
| 4V38 | C | 0.7486 | 1.1010 | 1.3918 | **14.1572** | 2.0860 |
| 5G1M | 89A | **5.6242** | **4.8734** | **6.1938** | **4.6805** | 3.9778 |
| 5G1M | NAG | 1.8182 | 2.5056 | 2.2855 | 2.1843 | 0.9695 |
| 5H9A | HVD | 1.4543 | 1.0974 | 1.1121 | 1.3381 | 1.0916 |
| 5NIA | DJ3 | 2.7926 | 1.6971 | 1.3176 | **9.8872** | 0.9454 |
| 5NZM | UPG | 1.6212 | 1.6926 | 1.6163 | 0.7229 | 1.6455 |
| 5ZA4 | NGE | 1.0337 | 0.8393 | 2.5825 | **9.7973** | 2.0491 |
| 6HB0 | GZL | 0.8326 | 1.1705 | 3.5339 | **4.1967** | 1.2658 |
| 6YPK | AHK | 3.6396 | 3.8413 | 3.2281 | **4.0769** | 3.8050 |

**Supplementary Table 4** Statistical significance between ProMoBind and ProMoBind-Base for predicting binding affinity on PDBBind v2020 dataset using time-based split. The p-values were obtained using a two-sided, paired t-test comparing the performance of ProMoBind and ProMoBind-Base across 5 runs.

| Methods | RMSE ↓ | MAE ↓ | Pearson ↑ | Spearman ↑ |
|---|---|---|---|---|
| ProMoBind-Base | 1.290 (0.013) | 1.041 (0.020) | 0.725 (0.005) | 0.694 (0.008) |
| ProMoBind | 1.268 (0.009) | 1.020 (0.009) | 0.738 (0.005) | 0.707 (0.011) |
| **P-value** | 0.0418 | 0.0279 | 0.0229 | 0.1126 |

**Supplementary Table 5** Comparison of model performance in predicting binding affinity on PDBBind v2020 using sequence identity-based split. The best results are highlighted in bold, and the second-best results are underlined.

| Methods | RMSE ↓ | MAE ↓ | Pearson ↑ | Spearman ↑ |
|---|---|---|---|---|
| PSICHIC | 1.565 (0.060) | 1.232 (0.050) | 0.584 (0.040) | 0.568 (0.039) |
| ProMoBind-Base | 1.520 (0.031) | 1.233 (0.028) | 0.588 (0.031) | 0.585 (0.032) |
| ProMoBind | **1.481 (0.017)** | **1.172 (0.020)** | **0.610 (0.011)** | **0.605 (0.013)** |

**Supplementary Table 6** Statistical significance between ProMoBind and baselines for predicting binding affinity on PDBBind v2020 using sequence identity-based split. The p-values were obtained using a two-sided, paired t-test comparing the performance of ProMoBind and baselines across 5 runs.

| Methods | P-value | | | |
|---|---|---|---|---|
| | RMSE | MAE | Pearson | Spearman |
| PSICHIC | 0.0200 | 0.0486 | 0.0374 | 0.0131 |
| ProMoBind-Base | 0.0454 | 0.0103 | 0.0905 | 0.0336 |

**Supplementary Table 7** Performance comparison across protein families on PDBbind v2020 using time-based split. Only protein families with more than 10 entries are included. For each metric, the better result between ProMoBind and ProMoBind-Base is highlighted in bold.

| Protein Family | N of Entries | RMSE ↓ | | MAE ↓ | | Pearson ↑ | | Spearman ↑ | |
|---|---|---|---|---|---|---|---|---|---|
| | | ProMoBind-Base | ProMoBind | ProMoBind-Base | ProMoBind | ProMoBind-Base | ProMoBind | ProMoBind-Base | ProMoBind |
| Enzyme | 123 | 1.349 | **1.323** | 1.078 | **1.058** | 0.631 | **0.658** | 0.636 | **0.648** |
| Epigenetic Regulator | 35 | 1.078 | **0.860** | 0.836 | **0.705** | 0.771 | **0.859** | 0.790 | **0.851** |
| Cytosolic Protein | 27 | 1.515 | **1.473** | 1.347 | **1.326** | 0.541 | **0.546** | 0.526 | **0.529** |
| Transcription Factor | 16 | 1.085 | **1.012** | 0.935 | **0.863** | 0.608 | **0.652** | 0.611 | **0.665** |

**Supplementary Table 8** Comparison of training and inference times for predicting binding affinity on PDBBind v2020 dataset. The best results are highlighted in bold, while '-' indicates that the data is not applicable to the corresponding model.

| Methods | Min/Epoch | NumEpoch | TrainTime ↓ (min) | InferenceTime ↓ (s/mol) |
|---|---|---|---|---|
| Pafnucy | 44.1860 | 18 | 777.6728 | 0.179 |
| OnionNet | 5.3787 | 38 | 204.3924 | 0.004 |
| IGN | 0.4946 | 51 | 25.4242 | 0.006 |
| SIGN | 2.4466 | 64 | 155.6035 | 0.011 |
| SMINA | - | - | - | 57.314 |
| GNINA | - | - | - | 13.139 |
| dMaSIF | 3.4297 | 23 | 79.5699 | 0.231 |
| TankBind | 14.1787 | 135 | 1908.4554 | 0.652 |
| GraphDTA | 2.1363 | 74 | 158.0828 | 0.003 |
| TransformerCPI | 10.1625 | 47 | 481.7004 | 0.010 |
| MolTrans | 2.9268 | 48 | 139.9009 | 0.027 |
| DrugBAN | 2.2342 | 72 | 160.8606 | 0.004 |
| DGraphDTA | 0.2558 | 90 | **22.9219** | 0.002 |
| WGNN-DTA | 2.5493 | 46 | 118.2867 | 0.004 |
| STAMP-DPI | 6.5604 | 46 | 299.1562 | 0.011 |
| PSICHIC | 3.5100 | 12 | 42.8220 | 0.008 |
| **ProMoBind** | 0.3020 | 132 | 39.8046 | **0.001** |

**Supplementary Table 9** Statistical significance between ProMoBind and baselines for predicting $k_{\text{off}}$ using random scaffold split. The p-values were obtained using a two-sided, paired t-test comparing the performance of ProMoBind and baselines across 10 runs.

| Methods | Source of Input | P-value | | | |
|---|---|---|---|---|---|
| | | RMSE | MAE | Pearson | Spearman |
| Random Forest | ESM-2 + ECFP4 | 0.0291 | 0.6832 | 0.0092 | 0.0219 |
| | ESM-2 + Uni-Mol | 0.0002 | 0.0014 | 0.0006 | 0.0068 |
| XGBoost | ESM-2 + ECFP4 | 0.0226 | 0.7067 | 0.0027 | 0.0333 |
| | ESM-2 + Uni-Mol | 0.0067 | 0.0173 | 0.0065 | 0.0305 |
| ProMoBind-Base | ESM-2 + Uni-Mol | 0.0051 | 0.0014 | 0.0060 | 0.0120 |

**Supplementary Table 10** Statistical significance between ProMoBind and baselines for predicting $k_{\text{off}}$ using sequence identity-based split. The p-values were obtained using a two-sided, paired t-test comparing the performance of ProMoBind and baselines across 5 runs.

| Methods | Source of Input | P-value | | | |
|---|---|---|---|---|---|
| | | RMSE | MAE | Pearson | Spearman |
| Random Forest | ESM-2 + ECFP4 | 0.0279 | 0.0020 | 0.0222 | 0.0162 |
| | ESM-2 + Uni-Mol | 0.0343 | 0.0011 | 0.0405 | 0.0327 |
| XGBoost | ESM-2 + ECFP4 | 0.0487 | 0.0247 | 0.0193 | 0.0379 |
| | ESM-2 + Uni-Mol | 0.0310 | 0.0350 | 0.0253 | 0.0044 |
| ProMoBind-Base | ESM-2 + Uni-Mol | 0.0259 | 0.0284 | 0.0119 | 0.0285 |

**Supplementary Table 11** Hyperparameter configurations for ProMoSite and ProMoBind across different prediction tasks. 'BS' stands for binding site, and '-' indicates that the hyperparameter is not applicable to the task.

| **Hyperparameter** | **ProMoSite** | | **ProMoBind** | |
|---|---|---|---|---|
| | General BS | Cryptic BS | Affinity | Kinetics |
| Number of layers in interaction module | 2 | 2 | - | - |
| $h$ in interaction module | 4 | 4 | - | - |
| $d$ in interaction module | 32 | 32 | - | - |
| $\lambda$ in pooling module | - | 0.1 | - | - |
| $t$ in clustering and ranking | 0.2 | 0.2 | - | - |
| $c$ in clustering and ranking | 50 | 50 | - | - |
| Dropout rate | 0.25 | 0.1 | 0.0 | 0.2 |
| $d'$ in fusion | - | - | 1024 | 1024 |
| Learning rate | $5 \cdot 10^{-5}$ | $1 \cdot 10^{-6}$ | $5 \cdot 10^{-5}$ | $5 \cdot 10^{-6}$ |
| Batch size | 8 | 8 | 256 | 4 |
| Max. number of epochs | 10 | 40 | 150 | 500 |

**Supplementary Table 12** Details of dataset statistics, data splits, and module direct outputs for each experiment.

| **Module Involved** | **Experiment** | **Dataset & Protein-ligand Pair Count** | **Data Split Approach** | **Module Direct Output** |
|---|---|---|---|---|
| ProMoSite | General Binding Site Prediction | Training & Validation:<br>scPDB$_{\text{COACH420}}$, 17,101<br>Test:<br>COACH420, 359 | Data split follows the approach in [42]:<br>Proteins from the scPDB v.2017 dataset that have either sequence identity > 50%, or ligand similarity > 0.9 and sequence identity > 30% to any structure in the corresponding test set, are removed to construct scPDB$_{\text{COACH420}}$ and scPDB$_{\text{HOLO4K}}$.<br>Each resulting scPDB subset is further split into 10 cross-validation folds based on their Uniprot IDs to avoid data leakage, with 9 folds used for training and 1 fold for validation. | The binding site annotation (Yes/No) of each residue with respect to a given ligand. |
| | | Training & Validation:<br>scPDB$_{\text{HOLO4K}}$, 10,626<br>Test:<br>HOLO4K, 7,847 | | The binding site annotation (Yes/No) of each residue with respect to a given ligand. |
| | Cryptic Binding Site Prediction | Training:<br>scPDB$_{\text{PocketMiner}}$, 19,254<br>Validation / Test:<br>PocketMiner, 28 / 38 | The same validation and test sets as in the previous study [14], curated based on PDB, are used.<br>Proteins from the scPDB v.2017 dataset that have sequence identity > 30% to any protein in the validation and test sets are removed to get scPDB$_{\text{PocketMiner}}$. | The binding site annotation (Yes/No) of each residue with respect to a given ligand. |
| | Pre-training for Integration into ProMoBind and Downstream Tasks | Training:<br>scPDB$_{\text{Pretrain}}$, 20,403<br>(6,878,337 residue labels)<br>Validation:<br>PocketMiner, 28 | The full scPDB v.2017 dataset is filtered by BioLiP to pre-train ProMoSite, which is subsequently used in ProMoBind. The PocketMiner validation set is used to select the best-performing model." | The binding site annotation (Yes/No) of each residue with respect to a given ligand. |
| ProMoBind | Binding Affinity Prediction | Training / Validation / Test: PDBbind v2020, 17,717 / 961 / 363 | Following prior studies [9, 21], a time-based split is used:<br>Data published before 2019 for training and validation, and data published in or after 2019 for testing. | The binding affinity of each protein-ligand pair. |
| | | Training / Validation / Test: PDBbind v2020, 15,454 / 849 / 363 | Based on the time-based split in [9, 21], proteins in the training and validation sets with sequence identity > 80% to any protein in the test set are removed. | The binding affinity of each protein-ligand pair. |
| | Binding Kinetics Prediction | Training, Validation & Test: Own $k_{off}$ dataset based on PDBbind-koff-2020 and BindingDB, 683 | A random scaffold split with an 8:1:1 ratio for the training, validation, and test sets is used, ensuring that the test set contains ligands with distinct scaffolds not present in the training or validation sets. | The binding kinetics ($k_{off}$) of each protein-ligand pair. |
| | | Training:<br>Own $k_{off}$ dataset, 520<br>Validation & Test:<br>Own $k_{off}$ dataset, 163 | All protein sequences in our curated $k_{off}$ dataset are clustered to ensure that no protein pairs from different clusters share > 80% sequence identity. The largest cluster (520 samples) is used for training, and the remaining clusters are randomly split with a 1:1 ratio as validation and test sets. | The binding kinetics ($k_{off}$) of each protein-ligand pair. |

**Supplementary Table 13** Distribution of major protein families for each dataset in each experiment.

| Experiments | Datasets | Protein Families (ChEMBL Level 1) | | | | | | | | | | | | | |
|---|---|---|---|---|---|---|---|---|---|---|---|---|---|---|---|
| | | Enzyme | Epigenetic Regulator | Transcrip-tion Factor | Cytosolic Protein | Ion Channel | Nuclear Protein | Membrane Receptor | Secreted Protein | Auxiliary Transport Protein | Surface Antigen | Transporter | Adhesion | Structural Protein | Other |
| General Binding Site Prediction | $\text{scPDB}_{\text{COACH420}}$ | 37.04% | 1.57% | 2.30% | 1.01% | 0.52% | 0.13% | 0.74% | 0.61% | 0.08% | 0.01% | 0.13% | 0.00% | 0.40% | 55.46% |
| | COACH420 | 32.30% | 0.00% | 2.06% | 0.69% | 0.00% | 0.00% | 0.00% | 0.69% | 0.69% | 0.00% | 0.00% | 0.00% | 0.00% | 63.57% |
| | $\text{scPDB}_{\text{HOLO4K}}$ | 30.06% | 2.34% | 0.63% | 0.41% | 0.43% | 0.19% | 1.12% | 0.24% | 0.03% | 0.02% | 0.19% | 0.00% | 0.07% | 64.27% |
| | HOLO4K | 36.24% | 0.26% | 1.47% | 1.49% | 1.31% | 0.22% | 0.03% | 0.98% | 0.46% | 0.00% | 0.00% | 0.00% | 0.10% | 57.45% |
| Cryptic Binding Site Prediction | $\text{scPDB}_{\text{PocketMiner}}$ | 41.18% | 1.23% | 1.89% | 1.58% | 0.66% | 0.11% | 0.55% | 0.53% | 0.07% | 0.01% | 0.12% | 0.05% | 0.36% | 51.65% |
| | PocketMiner | 30.30% | 3.03% | 0.00% | 1.52% | 0.00% | 0.00% | 0.00% | 0.00% | 0.00% | 0.00% | 0.00% | 0.00% | 0.00% | 65.15% |
| Pre-training for Downstream Tasks | $\text{scPDB}_{\text{Pretrain}}$ | 41.55% | 1.36% | 2.26% | 1.51% | 0.65% | 0.11% | 0.65% | 0.47% | 0.11% | 0.01% | 0.11% | 0.04% | 0.34% | 50.84% |
| Binding Affinity Prediction | $\text{PDBbind}_{\text{train+valid}}$ (time-based) | 53.77% | 4.59% | 3.07% | 2.60% | 1.35% | 0.85% | 0.79% | 0.78% | 0.45% | 0.34% | 0.17% | 0.09% | 0.03% | 31.12% |
| | $\text{PDBbind}_{\text{train+valid}}$ (sequence identity-based) | 54.13% | 3.59% | 3.29% | 1.50% | 1.43% | 0.93% | 0.69% | 0.85% | 0.51% | 0.13% | 0.20% | 0.10% | 0.04% | 32.60% |
| | $\text{PDBbind}_{\text{test}}$ | 33.88% | 9.64% | 4.41% | 7.44% | 0.28% | 1.65% | 1.38% | 1.38% | 0.00% | 0.55% | 0.00% | 0.00% | 0.00% | 39.39% |
| Binding Kinetics Prediction | Own $k_{off}$ dataset | 72.18% | 1.76% | 0.73% | 1.32% | 1.32% | 1.90% | 1.32% | 1.02% | 0.00% | 0.00% | 0.00% | 0.00% | 0.15% | 18.30% |

### Supplementary Figures

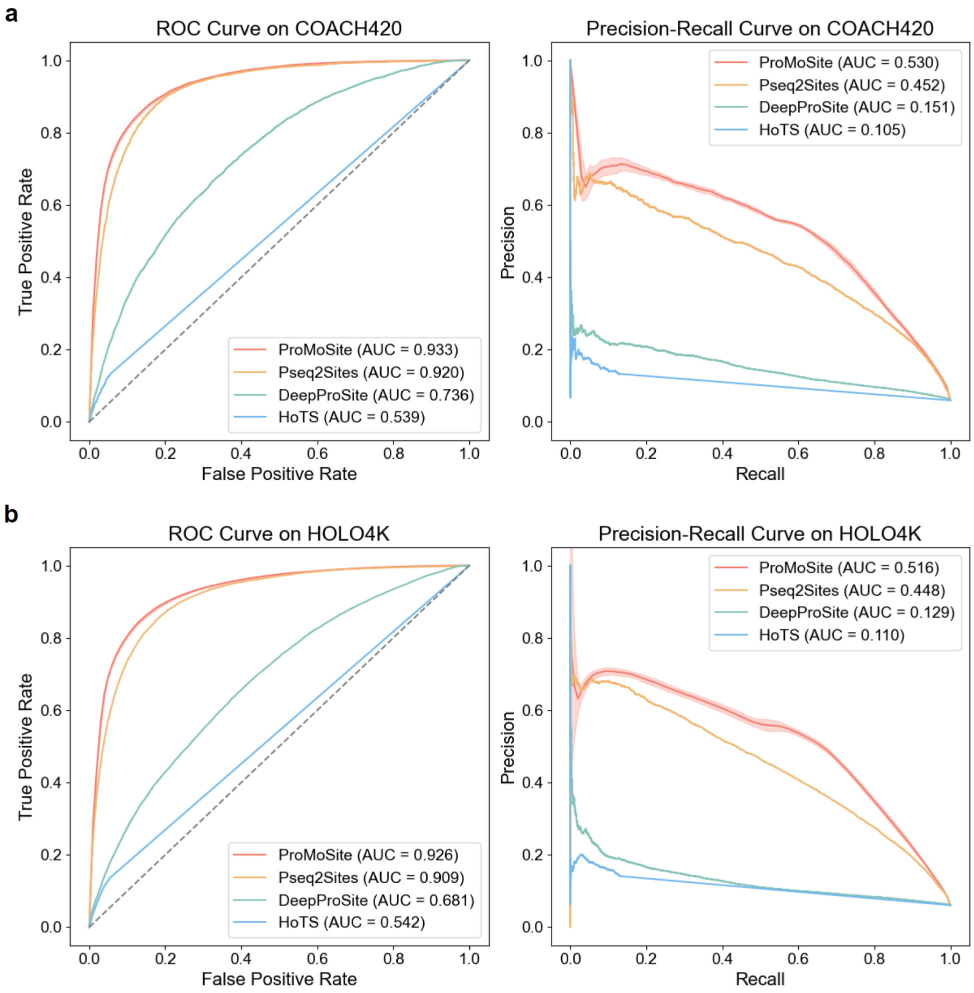


**Supplementary Fig. 1** Performance comparison of ProMoSite and sequence-based baselines on general binding site prediction. (**a**) ROC curves and Precision-Recall curves on COACH420 dataset. (**b**) ROC curves and Precision-Recall curves on HOLO4K dataset. ProMoSite consistently outperforms baselines in both datasets, demonstrating its superior predictive capability.

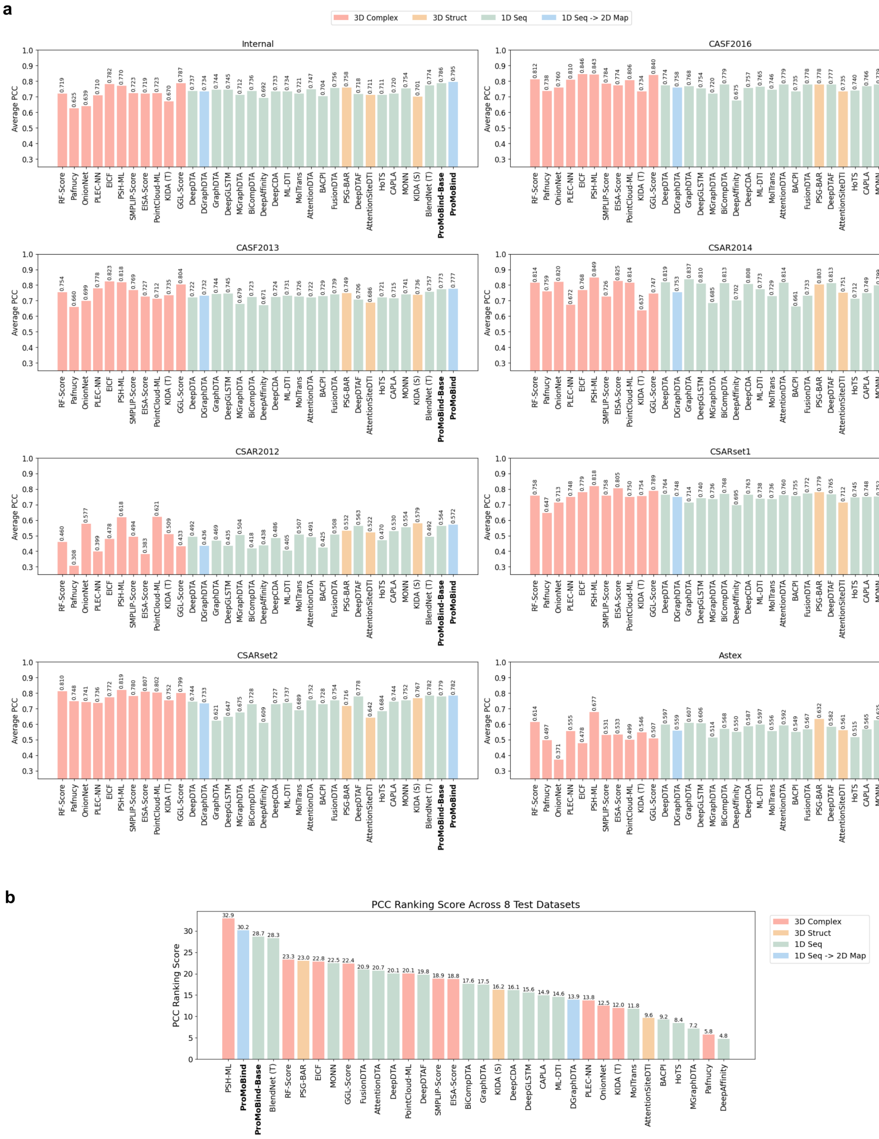


**Supplementary Fig. 2** Comparison of binding affinity prediction performance. (**a**) Model performance on eight test datasets from multiple sources, each containing experimentally determined protein–ligand complexes. Performance was quantified using the Pearson correlation coefficient (PCC), averaged across five-fold cross-validation. (**b**) Model ranking scores based on PCC averaged across the eight test datasets. A higher ranking score indicates better overall performance.

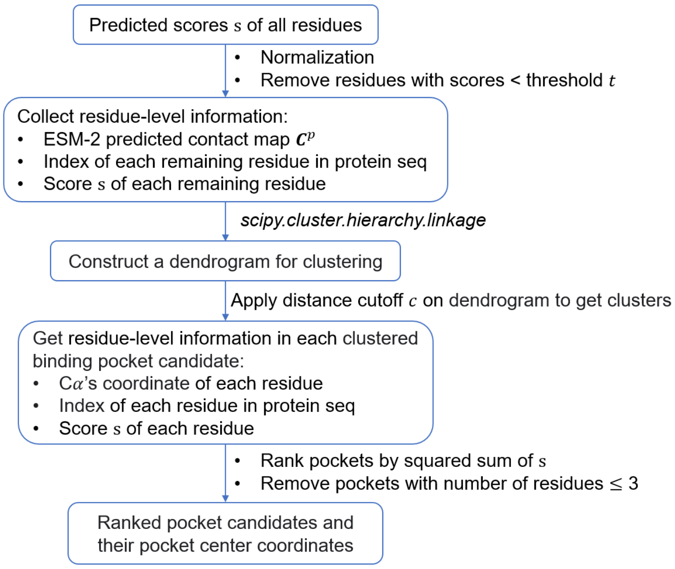


**Supplementary Fig. 3** Illustration of the clustering and ranking steps in ProMoSite.
